# Risky choice: probability weighting explains Independence Axiom violations in monkeys

**DOI:** 10.1101/2021.11.11.468261

**Authors:** Simone Ferrari-Toniolo, Leo Chi U Seak, Wolfram Schultz

## Abstract

Expected Utility Theory (EUT) provides axioms for maximizing utility in risky choice. The Independence Axiom (IA) is its most demanding axiom: preferences between two options should not change when altering both options equally by mixing them with a common gamble. We tested common consequence (CC) and common ratio (CR) violations of the IA over several months in thousands of stochastic choices using a large variety of binary option sets. Three monkeys showed consistently few outright *Preference Reversals* (8%) but substantial graded *Preference Changes* (46%) between the initial preferred gamble and the corresponding altered gamble. Linear Discriminant Analysis (LDA) indicated that gamble probabilities predicted most *Preference Changes* in CC (72%) and CR (88%) tests. The Akaike Information Criterion indicated that probability weighting within Cumulative Prospect Theory (CPT) explained choices better than models using Expected Value (EV) or EUT. Fitting by utility and probability weighting functions of CPT resulted in nonlinear and non-parallel indifference curves (IC) in the Marschak-Machina triangle and suggested IA non-compliance of models using EV or EUT. Indeed, CPT models predicted *Preference Changes* better than EV and EUT models. Indifference points in out-of-sample tests were closer to CPT-estimated ICs than EV and EUT ICs. Finally, while the few outright *Preference Reversals* may reflect the long experience of our monkeys, their more graded *Preference Changes* corresponded to those reported for humans. In benefitting from the wide testing possibilities in monkeys, our stringent axiomatic tests contribute critical information about risky decision-making and serve as basis for investigating neuronal decision mechanisms.

## Introduction

Most decisions we face include some degree of uncertainty. Economic decision theories that quantify the uncertainty associated with the choice options propose rigorous mathematical foundations for choice under risk. Expected Utility Theory (EUT), large parts of which were formalized by von Neumann and Morgenstern (1944), defines a mathematical framework based on four simple axioms, completeness, transitivity, continuity and independence. These axioms constitute the necessary and sufficient conditions for maximizing a specific subjective quantity, Expected Utility (EU): we simply choose the option with the highest EU. Utility (*U*) is the subjectively assigned value to a reward magnitude *m* (*U* = *u*(*m*)), and the subjective value of a probabilistic reward corresponds to the expected value of the utility distribution, called Expected Utility (*EU* = ∑ *u*(*m*_*i*_) · *p*_*i*_).

The independence axiom (IA) constitutes the fourth EUT axiom and is central to defining EU as subjective value. Together with the continuity axiom, the IA defines how magnitude and probability are combined to compute the global, subjective value of a risky choice option. The IA had been implicitly assumed by von Neumann and Morgenstern in their description of EUT tests and formulated, discussed and empirically tested by Marschak (1950), Allais (1953) and Savage (1954). The IA states that our preferences should not change when mixing all choice options with a common gamble. However, experiments have shown for decades that humans fail to comply with the IA (Allais, 1953; Kahneman & Tversky, 1979; Loomes & Sugden, 1987; Moscati, 2016; Starmer, 2000), which motivated additions to the existing utility theories, including prominently Prospect Theory (Kahneman & Tversky, 1979; Tversky & Kahneman, 1992).

Several sources of IA violations have been proposed, including subjective probability weighting, the certainty effect, the fanning-out hypothesis and heuristic schemes (Camerer, 1989; Kahneman & Tversky, 1979; Katsikopoulos, Gigerenzer, Katsikopoulos, & Gigerenzer, 2008; Machina, 1982; Savage, 1954). Past studies have also suggested that violations may not be as systematic as initially thought, reporting a significant proportion of EUT-compliant subjects (Harless & Camerer, 1994; Hey & Orme, 1994). Moreover, among the studies showing significant failures of the IA, high variability and conflicting types of violations have been reported (Battalio, Kagel, & Jiranyakul, 1990; P. Blavatskyy, Ortmann, & Panchenko, 2022; Conlisk, 1989; List & Haigh, 2005; Ruggeri et al., 2020; G. Wu & Gonzalez, 1998). The type and strength of violations also differed among distinct populations of subjects (Huck & Müller, 2012). Finally, human choices were usually tested with a small choice set and not repeated, missing effects of choice variability within each subject. Altogether, these results leave a fragmented picture on the extent, types and causes of IA violations. Clarifying these aspects would crucially contribute to understanding the mechanism underlying economic decisions.

The IA has not been tested in non-human primates, leaving an open question about the limits of compliance with EUT of our closest, experimentally viable, evolutionary relative. Monkeys can choose between actual outcomes that are tangibly delivered after every choice (as opposed to hypothetical outcomes) and can perform hundreds of daily choices. Monkeys allow systematic and incentive-compatible tests of EUT axioms in the same subject across a wide range of tests. Their reliable and stable performance minimizes errors and rules out insufficient learning, as noted for rodent tests of the IA axiom (Camerer, 1989; Kagel, Macdonald, Battalio, Kagel, & Mac, 1990). Monkeys’ choices satisfy first-, second- and third-order stochastic dominance, allow comparisons between risky and riskless utility functions, reveal nonlinear probability weighting, comply with the EUT continuity axiom, and can respect the Independence of Irrelevant Alternatives of two-component bundles (Bujold, Seak, Schultz, & Ferrari-Toniolo, 2021; Ferrari-Toniolo, Bujold, Grabenhorst, Báez-Mendoza, & Schultz, 2021; Ferrari-Toniolo, Bujold, & Schultz, 2019; Genest, Stauffer, & Schultz, 2016; Pastor-Bernier, Plott, & Schultz, 2017; Pelé, Broihanne, Thierry, Call, & Dufour, 2014; Stauffer, Lak, Bossaerts, & Schultz, 2015; Stauffer, Lak, & Schultz, 2014). However, without testing the IA, these choice data do not yet allow us to identify specific forms of subjective value computation. Work on monkeys is particularly suitable for achieving this goal, as the typical collection of large data sets facilitates thorough comparisons of economic models. As ultimate goal, well-defined behavioral assessments of EUT axioms, and in particular of the IA, would allow stringent, concept-based brain investigations of economic choice mechanisms with the high precision of primate single-cell neurophysiology. Given the evolutionary relationship between humans and monkeys, evidence of similar IA violations in the two species would help further our understanding of human decision making, both from an economic perspective and from a neurophysiological one.

Here, we used the IA to test the conditions and limits of utility-maximizing stochastic choices in three rhesus monkeys. The animals performed thousands of choices between gambles to identify specific forms of value computation, notably utility and probability-weighting. We systematically varied the gambles’ probabilities in common consequence and common ratio tests across the whole Marschak-Machina triangle to gain a comprehensive and detailed view of axiom compliance and violation. The animals consistently showed relatively few outright *Preference Reversals*, possibly due to their extended experience with the gambles, but substantial graded *Preference Changes*. Comparisons between economic model fits to the measured choices demonstrated that the probability weighting of Cumulative Prospect Theory (CPT) explained the choices better than models using Expected Value (EV) or EUT. The graded *Preference Changes* in our monkeys compared in frequency and strength to those reported for humans. These axiom-driven experiments identified the critical decision variables for utility-maximizing choices according to the IA and provide a basis for investigating the underlying neuronal signals in primates.

## Methods

### Animals

Three adult male rhesus macaques (*Macaca mulatta*) were used in this experiment: Monkey A (13 kg), Monkey B (11.5 kg) and Monkey C (11 kg). The animals were born in captivity at the Medical Research Council’s Centre for Macaques (CFM) in the UK. Monkey A (‘Tigger’) and Monkey B (‘Ugo’) had been surgically implanted with a headpost and a recording chamber for neurophysiological recording; they were headposted for 2 - 3 hours on each test day of the current experiment, which was intermingled with neuronal recordings on separate days. Both animals had previous experience with the visual stimuli and experimental setup (Ferrari-Toniolo et al., 2019).

Monkey C (‘Aragorn’) had no implant, no head posting and no previous task experience. All experimental procedures had been ethically reviewed and approved and were regulated and continuously supervised by the following institutions and individuals in the UK and at the University of Cambridge (UCam): the Minister of State at the UK Home Office, the Animals in Science Regulation Unit (ASRU) of the UK Home Office implementing the Animals (Scientific Procedures) Act 1986 with Amendment Regulations 2012, the UK Animals in Science Committee (ASC), the local UK Home Office Inspector, the UK National Centre for Replacement, Refinement and Reduction of Animal Experiments (NC3Rs), the UCam Animal Welfare and Ethical Review Body (AWERB), the UCam Governance and Strategy Committee, the Home Office Establishment License Holder of the UCam Biomedical Service (UBS), the UBS Director for Governance and Welfare, the UBS Named Information and Compliance Support Officer, the UBS Named Veterinary Surgeon (NVS), and the UBS Named Animal Care and Welfare Officer (NACWO).

### Task design

Each animal was seated in custom-made a primate chair (Crist instruments) in which he chose on each trial between two discrete and distinct options that were simultaneously presented at the right and left on a computer monitor at a distance of 50 cm in front of it. The animal indicated its choice by moving a joystick (Biotronix Workshop, University of Cambridge) either to the right or the left by an equal distance. The position of the joystick was monitored via custom code using Psychtoolbox 3 in Matlab (The MathWorks). The animals were first trained in >10,000 trials to learn the independently set reward magnitudes (*m*) and probabilities (*p*) that were indicated by a specifically set visual stimulus. Reward magnitude was signaled by the vertical position of a horizonal line; the probability of receiving that reward magnitude was proportional to the length of the horizonal line away from stimulus center (Fig. 1*A*). A stimulus with a full-length, single horizontal line corresponded to a sure reward (i.e. a degenerate gamble, *p* = 1), whereas multiple horizontal lines with less than full length indicated multiple possible gamble outcomes. At the end of each trial, the chosen option, and no other option, was paid out. From that paid-out option, one, and only one, of the outcomes was delivered to the animal. Thus, both the options and the outcomes of each option were mutually exclusive and collectively exhaustive. We used three fixed reward magnitudes: 0 ml (low; *m1*), 0.25 ml (middle; *m2*) and 0.5 ml (high; *m3*) of the same fruit juice or water; reward probabilities of the three reward magnitudes (*p1, p2, p3*) varied between 0 and 1, with a minimum step of 0.02. All option sets alternated pseudo-randomly. More details can be found in our previous study employing the same presentation design (Ferrari-Toniolo et al., 2021).

**Fig. 1.**
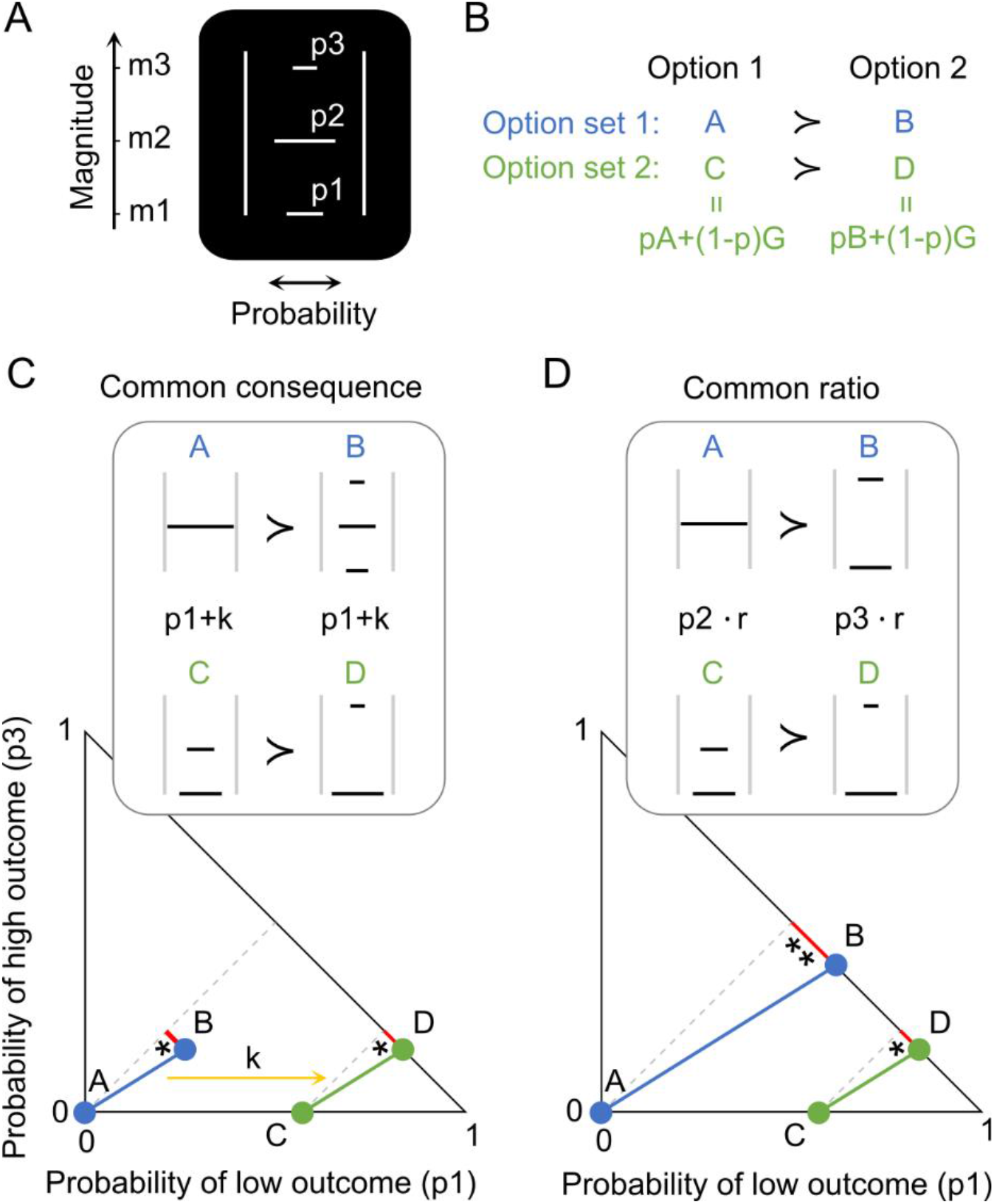
Experimental design for testing the independence axiom (IA). (A) Visual stimulus predicting a three-outcome gamble. The vertical position of each horizontal bar represents reward amount (*m1, m2, m3*; ml of juice); the length of each bar represents reward probability (*p1, p2, p3*) of the respective amounts *m1, m2, m3*. (B) Principle of testing the IA with two option sets {A,B} and {C,D}. Options C and D are obtained by adding the same gamble G to both options A and B, weighted by probability p. (C) Common consequence test and its representation in the Marschak-Machina triangle. The x-and y-axes represent the probabilities of the low outcome (*p1*) and high outcome (*p3*), respectively (probability of middle outcome: *p2 = 1-p1-p3*). Blue dots represent the original option set {A,B}, and green dots represent its modified set {C,D} for testing the IA. The yellow arrow indicates how option set {A,B} becomes option set {C,D} by adding the same probability k to the probability *p1* of the low outcome *m1* in both gambles A and B. Grey lines connect gambles with same expected value and highlight the linear and parallel nature of indifference curves tested by the IA. IA compliance requires same preferences: if (A ≻ B) then (C ≻ D), or if (A ≻ B) then (C ≻ D) and same choice probabilities. (D) Common ratio test. Multiplication of option set {A,B} by the same ratio r results in test option set {C,D}. Option set {A,B} becomes option set {C,D} by multiplying a common ratio *r* with the probability of the middle outcome (*p2*) of gamble A and with the probability of the high outcome (*p3*) of gamble B. * and ** and lengths of red lines indicate different distances of gambles from expected values (grey lines), which may result in potential preference changes between the two option sets without necessarily indicating IA violation.

### Revealed preference and choice indifference

Each animal’s preferences were considered to be revealed from its choices and used as a basis for quantification in later analyses. Preference was defined as the probability of choosing one gamble over the alternative gamble in the same binary option set; a gamble was considered to be revealed preferred to another gamble if the first gamble was chosen with *P* > 0.5. Thus, for a binary option set {A,B}, a stochastic preference relation was defined as *P*(A|{A,B}) = N_A_/N_AB_, where N_A_ was the number of trials in which the monkey chose A over B, and N_AB_ was the total number of trials with the {A,B} gamble set. When *P*(A|{A,B}) > 0.5 (i.e. when A was chosen in more than 50% of the trials), the monkey stochastically revealed preferred A to B. We used the binomial test (*P* < 0.05; 1-tailed) to assess the statistical significance of such preference relation in a specific direction (either *P*{A,B} > 0.5 or *P*{A,B} < 0.5) against choice indifference (probability of choosing each option with *P* = 0.5). When *P*(A|{A,B}) = 0.5, the animal was indifferent between two options A and B (i.e. gamble A was as much revealed preferred as gamble B).

### Defining the IA

The IA states that for any gamble A that is preferred to a gamble B, the combination of gamble A with gamble G should be preferred to the combination of gamble B with gamble G; the combined options are themselves gambles and are called C and D. Compliance with IA requires that the commonly added gamble G does not change the preference for the options.

Thus, individuals who prefer gambles A to B should also prefer gambles C to D (Fig. 1*B*). Any preference change constitutes an IA violation. These notions are formally stated as follows:

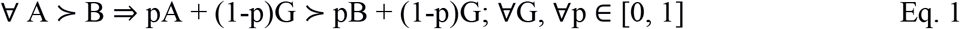

with A, B and G as gambles and p as probability. Gamble A was always a degenerate gamble with only a safe middle reward (*m2, p2* = 1), as used in Allais’ original test (Allais, 1953).

### Testing the IA

We assessed IA compliance in two commonly used tests: the common consequence (CC) test and the common ratio (CR) test.

The CC test consisted of adding (or subtracting) the same specific probability of an outcome (‘common consequence’) commonly to both options. Gamble A had a single outcome (*m2* = 0.25 ml, *p2* = 1), gamble B had three outcomes with fixed magnitudes *m1, m2, m3* and varying probabilities *p1, p2, p3* (Fig. 1*C*). For the CC test, we added a common probability *k* to the probability of the lower outcome (*p1*) to gambles A and B, which defined two new gambles C and D; probability *k* equals *p2* of option B. Adding probability *k* to the *p1* of gambles A and B consisted of reducing the original *p2* by *k* in both gambles (and thus reducing *p2* in gamble B to 0) to maintain the sum = 1.0 of all probabilities in each gamble.

The CR test consisted of multiplying the same ‘common ratio’ with the probabilities of all non-zero outcomes commonly for both gambles A and B. Gamble A had the same single outcome as in the CC test (*m2* = 0.25 ml, *p2* = 1), but gamble B had two outcomes with fixed magnitudes *m1, m3* and varying probabilities *p1, p3* (Fig. 1*D*). For the CR test, we multiplied a common ratio *r* with the probability of the middle outcome (*p2*) of gamble A and with the probability of the high outcome (*p3*) of gamble B and thus defined two new gambles C and D (thus, *p2* and *p3* of the new gambles C and D equalled *p2* and *p3* of gambles A and B commonly multiplied by *r*; *p*1’s of gambles C and D were adjusted to maintain the sum = 1.0 of all probabilities in each gamble).

The Marschak-Machina triangle constitutes an elegant scheme for graphically representing choice options and highlighting the predictions and consequences of IA compliance and violation (Machina, 1982; Marschak, 1950). In this abstract space (Fig. 1*C, D*, bottom), the x-axis represents the probability of obtaining the low outcome (*p1*) and the y-axis represents the probability of obtaining the high outcome (*p3*). The probability of the middle outcome (*p2*) derives from all probabilities summing to 1.0: *p2 =* 1 *- p1 - p3*. Each point inside the triangle represents a gamble. An IA test with two option sets is represented by two parallel lines that connect the original gambles A and B (blue), and the compounded gambles C and D (green) (note that these lines simply connect the gambles and do not indicate choice indifference; see below). In CC tests, the probability of the highest magnitude (*p3*) remains unchanged between gambles B and D (Fig. 1*C* bottom); thus, gamble D has the same vertical position as gamble B (y-axis). Thus, the change from option set {A,B} to option set {C,D} is graphically represented by a horizontal shift of the option set by probability k. In CR tests, probability *p3* changes between gambles B and D; these gambles have different vertical positions along the hypotenuse (Fig. 1*D* bottom). The change from option set {A,B} to option set {C,D} is graphically represented by a horizontal shift of gamble A to become gamble C, and by a downward shift along the hypotenuse from gamble B to gamble D.

In the Marschak-Machina triangle, equally revealed preferred gambles are connected by indifference curves (IC), whereas unequally preferred gambles are positioned on different ICs (see Fig. 5). While ICs should be linear and parallel with physical Expected Values (EV) and with utilities estimated according to EUT, they become non-parallel or curved with violations of the IA axiom.

### Definitions of IA violations

We used two measures for IA violation, *Preference Reversal* and *Preference Change*, both of which were based on the revealed preference of gamble A to gamble B indicated by the probability of choice of the initial gambles *P*(A|{A,B}). Thus, we used stochastic choices and correspondingly stochastic models to test the IA. While both outright *Preference Reversal* and graded significant *Preference Change* can detect IA violations, they cannot assess significant axiom compliance, as this corresponds to the null hypothesis of our statistical test.

Nonlinear and non-parallel indifference curves in the Marschak-Machina triangle demonstrate IA violations, as proposed by several non-EU theories (Bhatia & Loomes, 2017; Bordalo, 2012; Kahneman & Tversky, 1979; Machina, 1982; Savage, 1954).

Our first, binary measure for IA compliance was *Preference Reversal*. When the fixed gamble A was stochastically revealed preferred to gamble B, *Preference Reversal* was manifested as stochastically preferring gamble D to gamble C:

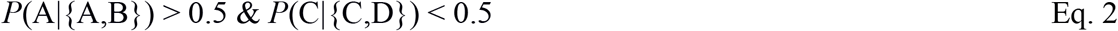

This reversal, in a non-stochastic setting, was originally observed in humans (Allais-type reversal; Allais 1953; Kahneman and Tversky 1979). To adapt this measure to stochastic choices, we assessed the significance of the initial preference *P*(A|{A,B}) > 0.5 in comparison with choice indifference (*P*(A|{A,B}) = 0.5) using the binomial test (statistical *P* < 0.05; 1-tailed). Then *Preference Reversal* was evidenced as significant stochastic preference in the opposite direction (*P*(C|{C,D}) < 0.5) (binomial test).

To the opposite, when gamble B was stochastically revealed preferred to gamble A, we defined *Preference Reversal* as stochastically preferring gamble C to gamble D:

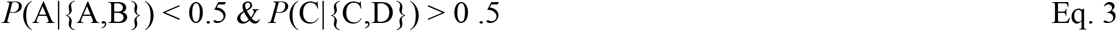

We assessed the significance of this reverse Allais-type *Preference Reversal* (P. R. Blavatskyy, 2013; Conlisk, 1989) in analogy to the (regular) Allais-type reversal.

Our second, more graded measure for IA violation was *Preference Change*. We used the metric S introduced by Conlisk (1989) who had assessed IA violations non-stochastically from single choices of multiple human participants. We adapted the Conlisk’S assessment to repeated, stochastic choices of individual animals and quantified *Preference Change* stochastically as ratio of probabilities of Allais-type and reverse Allais-type reversals:

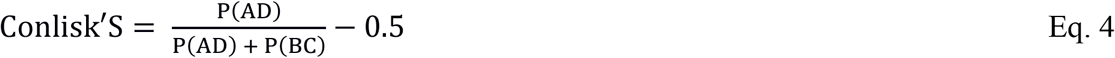

P(AD) indicates the probability of Allais-type reversals: *P*(A|{A,B}) > 0.5 and *P*(D|{C,D}) > 0.5. *P*(BC) indicates the probability of reverse Allais-type reversals: *P*(B|{A,B}) > 0.5 and *P*(C|{C,D}> 0.5. We set Conlisk’S = 0 when P(AD) + P(BC) = 0.

Assuming that choices in different trials were independent, we computed *P*(AD) = *P*(A|{A,B}) · *P*(D|{C,D}), obtaining:

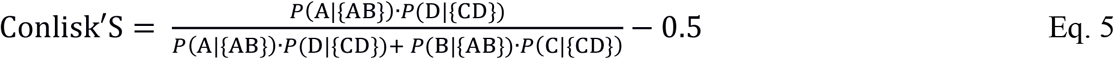

The Conlisk’S measure is a real number that varies between -0.5 and 0.5. We defined the significance of Conlisk’S as difference from zero (*P* < 0.05 on pooled sessions from a given monkey; one-sample t-test). To avoid unreasonably large violation measures from infrequent violations, we weighted the Conlisk’S with respect to the total proportion of violations and obtained the *Preference Change* S:

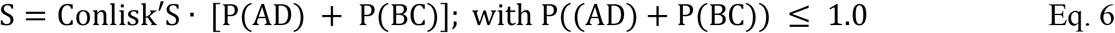

In this way, S is a real number that still varies between -0.5 and 0.5 and indicates the same direction of systematic IA violation as Conlisk’S but corresponds better to the fraction of trials producing the violation. The S is a measure of how much the preferences vary between option sets {A,B} and {C,D} (i.e. the non-vertical negative and positive slopes, indicating S > 0 and S < 0, respectively, in our preference comparisons in Figs. 2 and 3, red). All subsequent analyses used this S as measure of *Preference Change*.

**Fig. 2.**
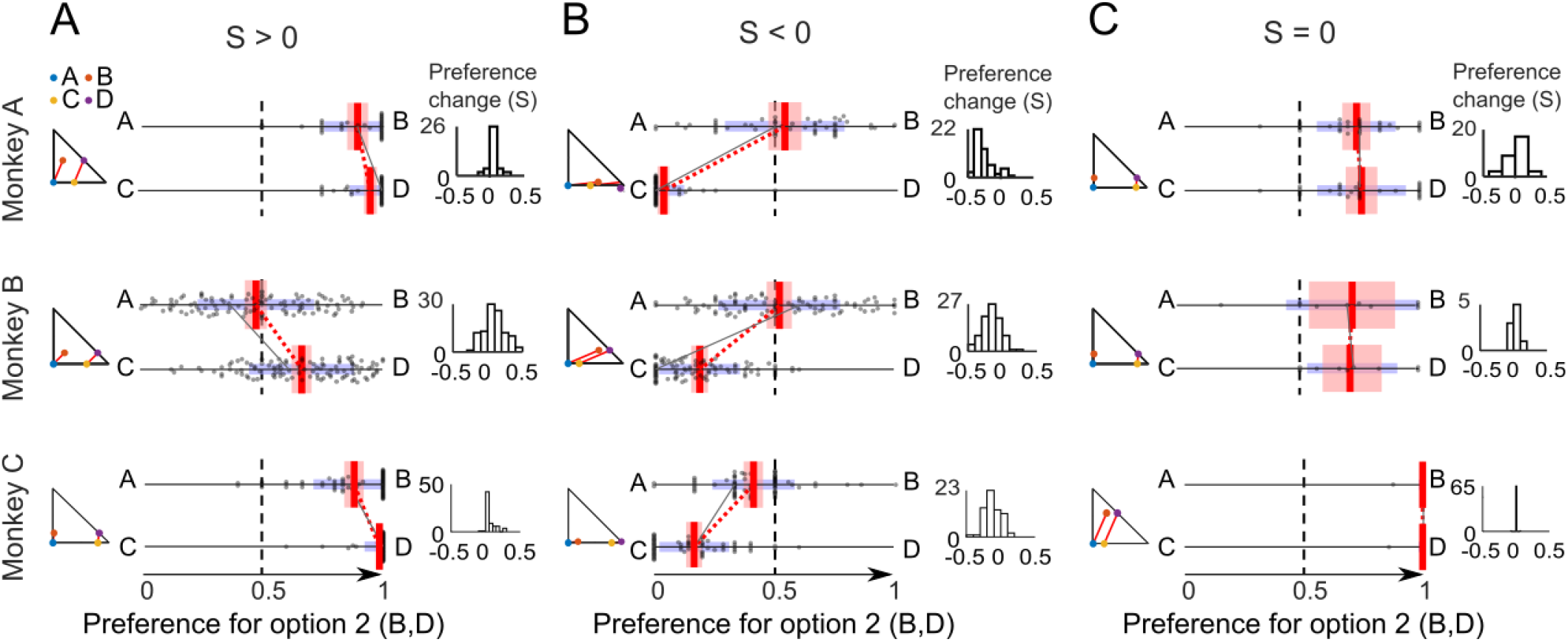
*Preference Changes* S during IA common consequence tests. (A) Significant violations with positive *Preference Change* measure S (*P* < 0.05 on pooled sessions from a given monkey; one-sample t-test against zero). Each panel shows at the left the small Marschak-Machina triangle for the tested options. The center plot shows the probability of choosing one option over the other option (Options A and C on the left; Options B and D on the right). Each dot represents the probability of choosing A over B or C over D in one session; red vertical bars represent averages of probability of choices (A - B or C - D) across sessions; red intervals show the 95% confidence interval; blue intervals show Standard Deviations (SD); the black line links preferences in one example session (red dotted line: average across sessions). Small histograms (right) show the distribution of S’s that quantifies the IA violation, across sessions. (B) Significant negative *Preference Changes* S. (C) Insignificant *Preference Changes* S.

**Fig. 3.**
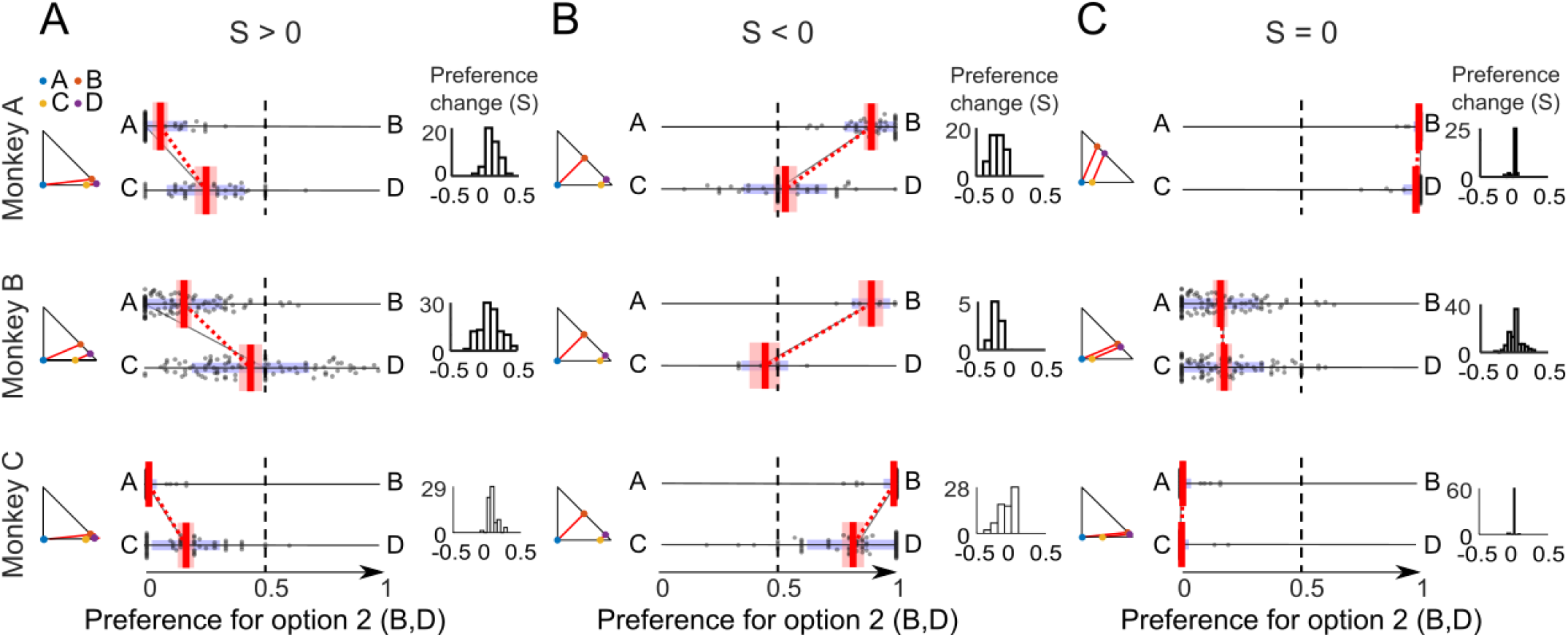
*Preference Changes* (S) during IA common ratio tests. (A) Significant positive *Preference Change* measure S (all *P* < 0.05; one-sample t-test). For conventions, see Fig. 2. (B) Significant negative *Preference Changes* S. (C) Insignificant *Preference Changes* S.

With repeated choices, IA violations may seem to occur simply because of some random variability in preferences, but the measure of S cancels out random violations in opposite directions and thus results in a robust measure of *Preference Change*. Compliance with the IA is manifested as *Preference Changes* resulting in S = 0, which reflects either an absence of IA violations or a balanced number of random IA violations in the two directions. *Preference Changes* are manifested by positive or negative S values that differ significantly from zero (*P* < 0.05; one-sample t-test).

In our stochastic version of EUT, *Preference Changes* (significant S ≠ 0) represent IA violations in CC tests in which gambles B and D have equal distance from the respective parallel choice indifference lines representing equal expected utility in the Marschak-Machina triangle (Fig. 1*C*); the equal utility difference should preserve their preference within their respective option sets and thus maintain linear and parallel ICs. By contrast, *Preference Changes* in CR tests are necessary but not sufficient for defining IA violations. The CR test places gambles B and D at different distances from the respective parallel indifference lines (Fig. 1*D*) that reflect different expected utility differences for the two option sets that may result in preference changes (but not outright preference reversals) without indicating IA violation. In other words, when using a stochastic model, linear and parallel ICs can produce non-zero S values in the CR test. Thus, non-zero S values do not necessarily indicate IA violations in CR tests.

### Classification analysis

To check whether the *Preference Changes* in the IA tests depended on reward probability, we performed classification analyses. We used a Linear Discriminant Analysis (LDA) classifier (*fitcdiscr* function in Matlab) to predict the sign of the *Preference Change* measure S. We characterized the changes with a leave-one-out procedure using Linear Discriminant Analysis (LDA). We trained the LDA with all data except for those from the predicted leave-out choice tests to build 36 models (18 CC tests and 18 CR tests) for each animal (we discarded one option set in CR tests with Monkey C in which S was zero). As each of the 36 models was used to predict the left-out data, we obtained 36 predictions for each animal. We compared these predictions to the measured directions of *Preference Change* to check the accuracy of the prediction. To illustrate the test sensitivity (true positive rate / ability to predict one class) and specificity (false positive rate / ability to predict the other class), we drew a confusion matrix for the CC and CR tests, separately for each animal (see Fig. 4).

**Fig. 4.**
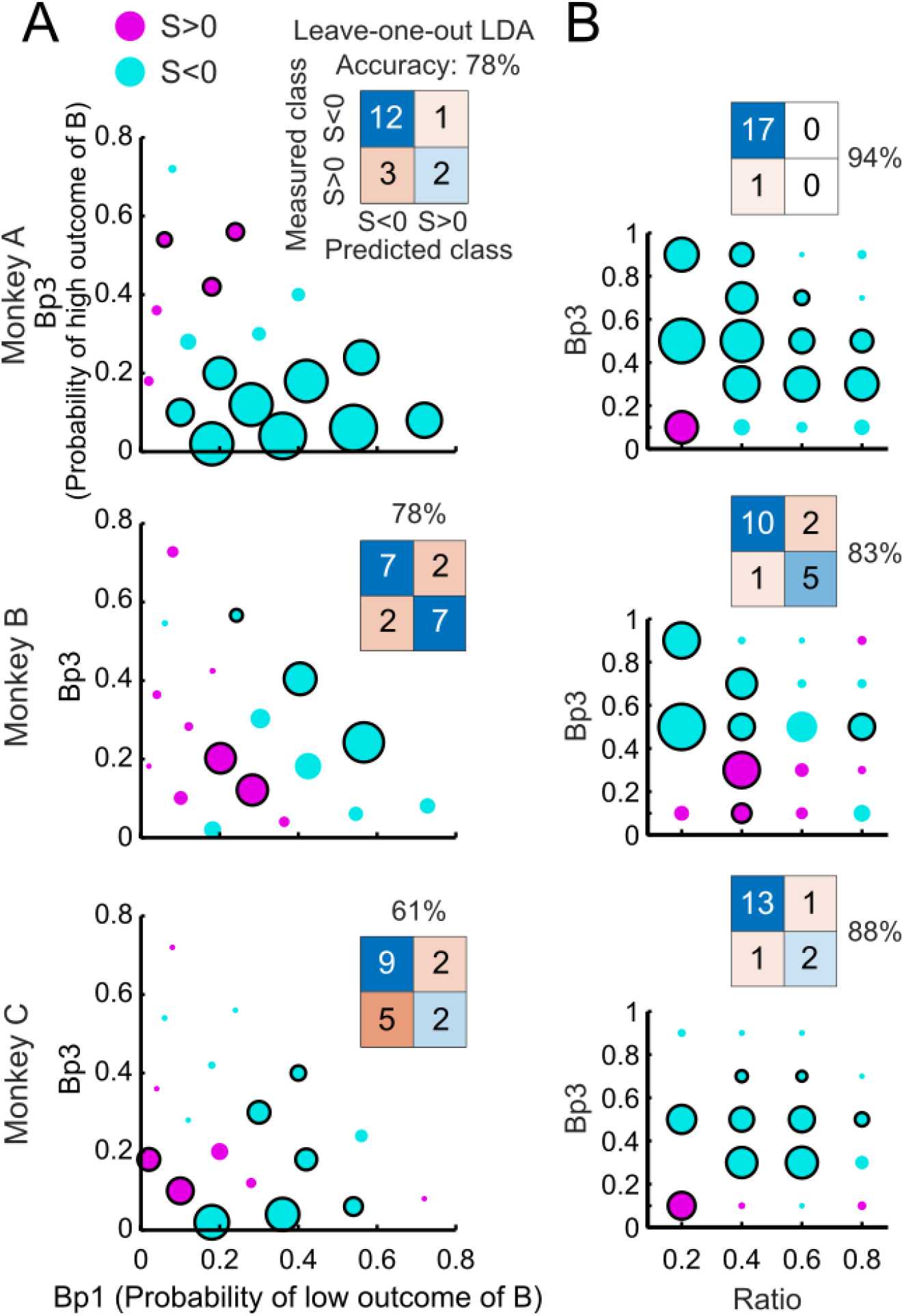
Probability dependency of IA *Preference Changes*. (A) Common consequence test. The x- and y-axes show the probabilities of low and high magnitudes of option B, respectively. Purple and cyan dots represent positive and negative *Preference Change* measures S, respectively. Black circles around dots indicate significance (*P* < 0.05; one-sample t-test). Insets show confusion matrices from classifications using Linear Discriminant Analysis (LDA). (B) Common ratio test. The x- and y-axes show the ration and the probability of high magnitudes of option B, respectively. One option set with S = 0 not shown in Monkey C.

**Fig. 5.**
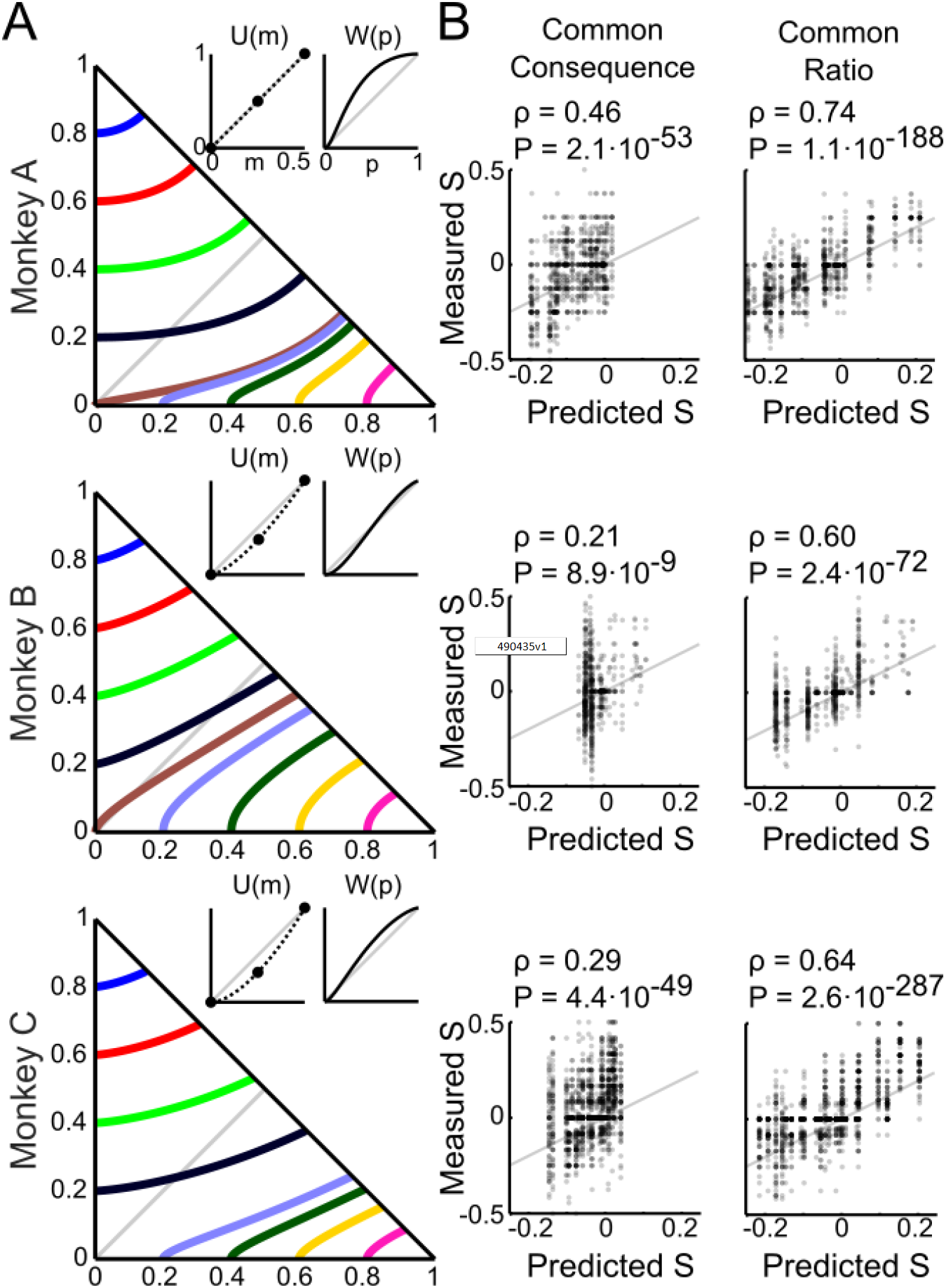
Cumulative prospect theory (CPT) modeling can explain the measured *Preference Changes*. (A) Choice indifference curves (ICs) in the Marschak-Machina triangle. Each line represents one IC, and all gambles on a given line are equally preferred to each other. Insets show estimated utility and probability weighting functions, using the CPT model, from all trials (common ratio and common consequence tests). The three monkeys had individually differing, mostly non-linear utility and probability weighting functions. Note that utility functions were only estimated for three points (black dots), which were the only three magnitudes used in the experiment (*m1, m2, m3*); thus, the dashed lines do not represent the full shape of the utility function. (B) Pearson correlations between measured S’s and S’s predicted by the CPT-modeled utility and probability weighting functions for the common consequence and common ratio tests.

### Economic modeling of choice behavior

We defined a standard discrete choice softmax function (McFadden, 2001) to describe stochastic preferences. The probability *P* of choosing a generic option A over another option B was defined as:

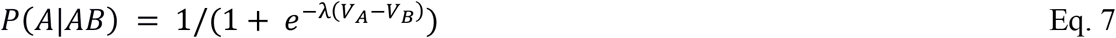

where λ represents the noise parameter, defining the steepness of the preference function (steeper for higher λ values). Based on EV theory, EUT and Cumulative Prospect Theory (CPT), we used three models to define the value (*V*) of gambles. These models returned different estimates of choice probability according to Eq. 7.

In the EV model, each option’s value was its objective *Expected Value*:

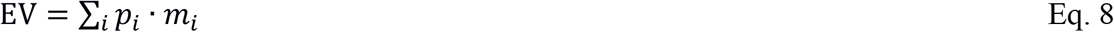

For a generic three outcome gamble in our task, it corresponded to (*m*_1_ was zero in our task and therefore *p*_1_ · *m*_1_ = 0):

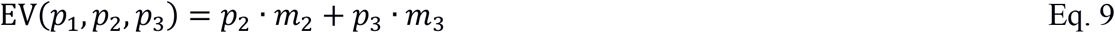

In the EUT model, each option’s subjective value was defined via the utility function (*u*) as its *Expected Utility*:

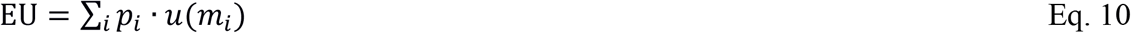

In our gambles’ space this mapped to:

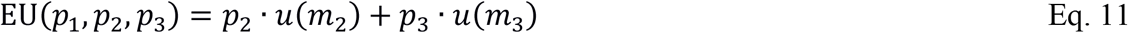

In the CPT model, each option’s subjective value was called a *Prospect Value* and defined by a utility function (*u*) together with a probability weighting function (*w*), combined in a cumulative form (Tversky & Kahneman, 1992):

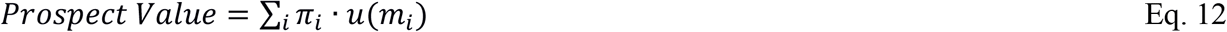

where *π*_*i*_ = *w* (*p*_*i*_ + … + *p*_*n*_) − *w* (*p*_*i*+1_ + … + *p*_*n*_), with *n* indicating the number of outcomes, and index *i* corresponding to the outcomes ordered from worst to best (*m*_1_ and *m*_3_ respectively, in our task). For a generic three-outcome gamble (with probabilities *p*_1_ *p*_2_ *p*_3_), Eq. 12 becomes:

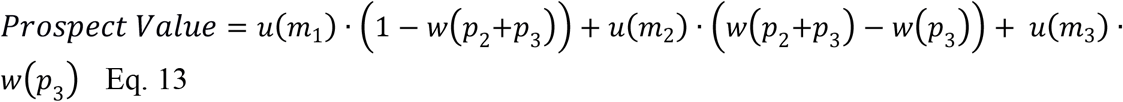

which, with our set of magnitudes and normalized utility, corresponds to

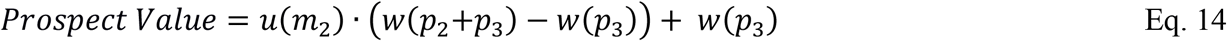

In these three value-estimating equations, each *p*_*i*_ represents the probability of getting the respective reward magnitude (*m*_*i*_): *p*_*i*_ and *m*_*i*_ represent the probability and magnitude of the lowest outcome (0 ml); *m*_2_ and *m*_2_ the probability and magnitude of the middle outcome (0.25 ml); *p*_3_ and *m*_3_ are relative to the highest outcome (0.5 ml). In the EUT and CPT models, the utility function was defined as a power function (free parameter ρ), normalized to the highest magnitude level:

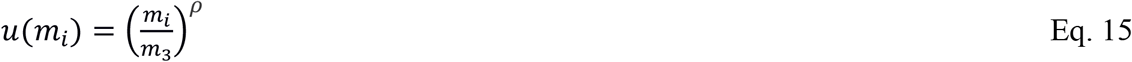

The ρ parameter defines a convex (ρ > 1) or concave (ρ < 1) utility function, with - = 1 corresponding to linear utility. Note that having only three magnitude levels in the current experiment implied that the only meaningful utility value was that of the middle outcome magnitude (*m2*) in relation to the other two outcomes (*m1* and *m3*). Thus, although a larger set of magnitudes may result in more complex utility functions, a power function would be sufficient to account for the difference in subjective evaluation of the three reward magnitudes used in our study.

In the CPT model, cumulative probability weighting was defined as a two-parameter *Prelec* function (Prelec, 1998; Stott, 2006) as in our earlier study (Ferrari-Toniolo et al., 2019):

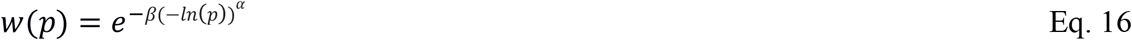

where *α* allows the function to vary from inverse-S-shaped (*α* < 1) to S-shaped (*α* > 1), while *β* shifts the function vertically.

We estimated the functions’ parameters (*θ*) with the maximum likelihood estimation (MLE) method, by maximizing the log-likelihood function defined (for a choice between generic options A and B; using *fminsearch* in Matlab) as:

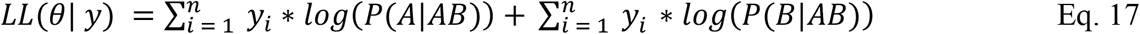

The experimental choice outcome was defined for each trial *i* by the binary variable *y*_*i*_ (1 when A chosen, 0 when B chosen) and 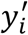 (1 when B chosen).

To validate our economic models, we used an out-of-sample dataset that consisted of a set of gambles that differed from the gamble set used for the IA tests. We presented monkeys with choices between one fixed option (J) on the x axis (*p1* between 0 and 0.8 in 0.2 increments) or on the y axis (*p3* between 0.2 and 0.8 in 0.2 increments) and another option (K) with variable *p1* (and *p3*) and fixed *p2* (with *p2* between 0.2 and 1, in 0.2 increments). For each option J, by varying the probability *p1* in option K, we identified an indifference point (IP) as the point within the triangle where a fitted softmax preference function would take the value of 0.5. All choice trials in the out-of-sample test were pseudo-randomly intermingled. IPs were estimated separately in each weekdaily session.

### Comparison with human choices

We tested whether the observed IA violations in the 18 CC tests in monkeys corresponded to the violations reported in 39 human studies (P. Blavatskyy, Ortmann, & Panchenko, 2015), using data pooled from all three monkeys. We used two different comparison methods, a confusion matrix using binary classes of *Preference Change* (either S > 0 or S < 0), and a Pearson correlation using real-number *Preference Change*s (S varying between -0.5 and +0.5).

However, the gambles used in our monkeys differed from the gambles used in the human studies (see Fig. 7*A*). Therefore, for more accurate comparisons, we first needed to predict the *Preference Changes* S that would have occurred in our monkeys had we used the exact same gambles as in humans; to control for directionality of testing, we also needed to predict, in the reverse direction, the *Preference Changes* S that would have occurred in the human studies had they used the same gambles as we did in our monkeys.

For the confusion matrix, we predicted the S’s for the unused gambles with an LDA classifier trained on the S’s of the actually used gambles. For predicting the monkey S’s for the gambles used in the human studies, we trained the LDA with the binary monkey S’s (S > 0, S < 0), and the probabilities for the low and the high magnitudes of the monkey gambles (*p1, p3)*. In the reverse direction, for predicting the human S’s for the gambles used in our monkeys, we trained the LDA with the binary human S’s, the probabilities for the low and the high magnitudes of the human gambles (*p1, p3*), and the ratio of the middle and high magnitudes of the human gambles (*m2* / *m3*). Then we used the confusion matrix to compare measured human S’s with predicted monkey S’s (see Fig. 7*C* left) and, vice versa, measured monkey S’s with predicted human S’s (see Fig. 7*D* left). The accuracy of the comparison was defined in percent from the ratio: (total number of successful comparisons) / (total number of comparisons). For example, in the confusion matrix shown in Fig. 7*C*, the total number of successful comparisons is (6 + 26) / (6 + 7 + 0 + 26) = 0.82, which equals 82%.

For the Pearson correlation, we predicted the S’s for the unused gambles with two different multiple linear regression systems depending on the direction of comparison. The regression for the comparison of measured human *Preference Changes* S with predicted monkey S’s first estimated the beta parameters for the S’s measured in monkeys as follows:

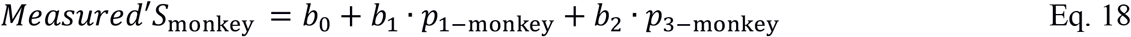

with p1-monkey and p3-monkey as probabilities of lowest and highest magnitude of the gambles used in option B in monkeys. Then we applied the estimated betas from Eq. 18 to all gambles used in the 39 human studies to predict the numeric *Preference Change* S for these gambles in monkeys:

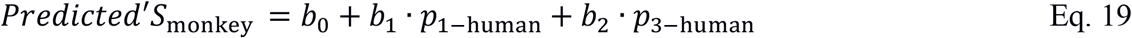

with p1-human and p3-human as probabilities of lowest and highest magnitude of the gambles used in option B in humans. Then we compared the measured human S’s with the predicted monkey S’s using a Pearson correlation (see Fig 7*C* right).

In the reverse direction, comparing measured monkey S’s with predicted human S’s, the regression first estimated the beta parameters for the *Preference Change* S measured in humans with the modified regression model:

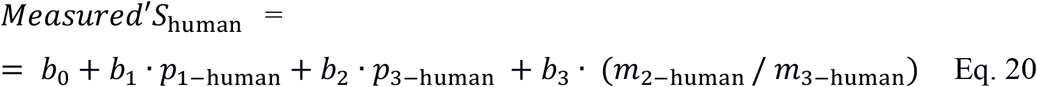

with p1-human and p3-human as probabilities of lowest and highest magnitudes of gamble B, and m2-human and m3-human as middle and highest magnitude used in humans (magnitudes varied across the human studies but were constant in all monkey gambles). Then we applied the estimated betas from Eq. 20 to all gambles used in our monkeys to predict the numeric *Preference Change* S for these gambles in humans:

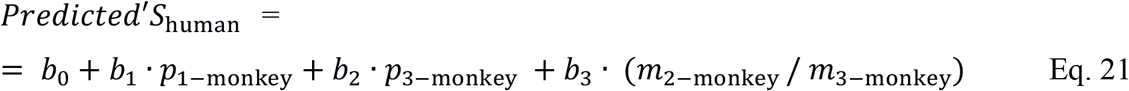

with p1-monkey and p3-monkey as probabilities of lowest and highest magnitudes of gamble B, and m2-monkey and m3-monkey as middle and highest magnitude used in monkeys. Then we compared the measured monkey S’s with the predicted human S’s using Pearson correlation (see Fig 7*D* right).

## Results

### Experimental design

We used stochastic choices to test compliance with the independence axiom (IA) in three monkeys. Two visual bar stimuli indicated two respective choice options on a computer monitor in front of the animal. The animal chose by moving a joystick towards one of the two options and 1.0 sec later received the reward of the chosen option. Each option was a gamble defined by three reward magnitudes (*m1, m2, m3*; ml of fruit juice) occurring with specific probabilities (*p1, p2, p3*; sum = 1.0) (Fig. 1*A*). Reward magnitude was indicated by bar height (higher was more), and the probability of delivering each magnitude was indicated by bar length away from stimulus center (longer was higher).

Testing the IA began with two gambles A and B that formed option set {A,B}. Gamble A was a degenerate gamble with safe and fixed middle reward magnitude (*m2* = 0.25 ml; *p2* = 1.0), whereas gambles B, C and D were two- or three-outcome gambles. The test gambles C and D derived from the common addition of gamble G and constituted option set {C,D} (Eq. 1).

Stochastic compliance with the IA requires that preferences do not change significantly between option sets {A,B} and {C,D} (Fig. 1*B*). We assessed the IA in the common consequence (CC) test and in the common ratio (CR) test (see Methods for definitions; Fig. 1*C, D*). When representing the gambles in the Marschak-Machina triangle, an IA test was plotted as a parallel shift of the line connecting the two gambles of each option set ({A,B} and {C,D}), with an additional line length change for a CR test.

### IA violations

We performed 18 different CC tests and 18 different CR tests in each of the three monkeys; each test was repeated on average 7.5 times per daily session for Monkey A, 17.7 times per session for Monkey B and 6.1 times per session for Monkey C. We systematically varied the reward probabilities and thereby tested the IA across the whole range represented by the Marschak-Machina triangle. We tested two violation directions: either gamble A was stochastically revealed preferred to gamble B (probability of choice *P*(A|{A,B}) > 0.5) and gamble D was stochastically preferred to gamble C (*P*(C|{C,D}) < 0.5) (Allais-type violation; Allais 1953), or gamble B was revealed preferred to gamble A (*P*(A|{A,B}) < 0.5) and gamble C was stochastically preferred to gamble D (*P*(C|{C,D}) > 0.5) (reverse Allais-type violation; Blavatsyy, 2013b). We considered two IA violation types, the more substantial binary *Preference Reversals* and the more subtle graded *Preference Changes*.

*Preference Reversals* across option sets {A,B} and {C,D} were defined by Eqs. 2 and 3 for AD and BC preference directions and tested for significance using the 1-tailed binomial test applied separately to Allais-type and reverse Allais-type reversals (*P* < 0.05; see Methods). IA violations indicated by significant *Preference Reversals* occurred only in a few of the 36 tests (N=8 for Monkey A, N=1 for Monkey B, N=0 for Monkey C; total of 8% (CC: 11%, CR: 6%); Table 1).

**Table 1.** *Preference Reversals* while testing the independence axiom.

|  | Test | *Preference Reversal* |  |
| --- | --- | --- | --- |
| Monkey<br>A | CC | $A > B \Rightarrow C < D$ | 0/18 |
| | | $A < B \Rightarrow C > D$ | 5/18 |
| | CR | $A > B \Rightarrow C < D$ | 0/18 |
| | | $A < B \Rightarrow C > D$ | 3/18 |
| Monkey<br>B | CC | $A > B \Rightarrow C < D$ | 1/18 |
| | | $A < B \Rightarrow C > D$ | 0/18 |
| | CR | $A > B \Rightarrow C < D$ | 0/18 |
| | | $A < B \Rightarrow C > D$ | 0/18 |
| Monkey<br>C | CC | $A > B \Rightarrow C < D$ | 0/18 |
| | | $A < B \Rightarrow C > D$ | 0/18 |
| | CR | $A > B \Rightarrow C < D$ | 0/18 |
| | | $A < B \Rightarrow C > D$ | 0/18 |

CC: common consequence test, CR: common ratio test. $x > y$ indicates 'x preferred to y', $x < y$ indicates 'y preferred to x'. $A > B$ leading to $C < D$ constitutes a stochastic Allais-type *Preference* *Reversal*, as indicated by significant probability of choice: $P(C|\{C,D\}) < 0.5$ (statistical $P < 0.05$ ; 1- tailed; binomial test) (see Eq. 2). By contrast, $A < B$ leading to $C > D$ constitutes a stochastic reverse Allais-type *Preference Reversal*: $(P(D|\{C,D\})) < 0.5$ (statistical $P < 0.05$ ) (see Eq. 3).

Note that all animals were highly familiar with gamble variations from tens of thousands of trials performed during several months of weekdaily experimentation.

*Preference Changes* were defined by Eqs. 4 - 6 that computed the variable S derived from Conlisk’S and tested for significance using a one-sample t-test against S = 0 in pooled sessions from a given monkey (*P* < 0.05). In contrast to the few outright *Preference Reversals*, significant *Preference Changes* using the metric S were rather frequent in all animals (N=21 for Monkey A, N=12 for Monkey B, N=17 for Monkey C; total of 46% (CC: 41%, CR: 52%); Table 2).

**Table 2.** *Preference Changes* while testing the independence axiom.

|  | Test | *Preference Change* |  |
| --- | --- | --- | --- |
| Monkey<br>A | CC | $S > 0$ | 1/18 |
| | | $S < 0$ | 8/18 |
| | CR | $S > 0$ | 1/18 |
| | | $S < 0$ | 11/18 |
| Monkey<br>B | CC | $S > 0$ | 2/18 |
| | | $S < 0$ | 3/18 |
| | CR | $S > 0$ | 2/18 |
| | | $S < 0$ | 5/18 |
| Monkey<br>C | CC | $S > 0$ | 2/18 |
| | | $S < 0$ | 6/18 |
| | CR | $S > 0$ | 1/18 |
| | | $S < 0$ | 8/18 |

CC: common consequence test, CR: common ratio test. *Preference Changes* were measured as real- number $S$ based on a modification of Conlisk's (see Eqs. 4 - 6) and tested for significance at $P <$ 0.05 (one-sample t-test against zero). *Preference Changes* represent IA violations for CC tests but are only necessary and not sufficient for claiming IA violations in CR tests.

*Preference Changes* are sufficient for defining IA violations in CC tests; here, gambles B and D have equal distance from the respective parallel choice indifference lines representing equal expected utility in the Marschak-Machina triangle (Fig. 1*C*); the equal utility difference should preserve their preference within their respective option sets and thus produce no violation.

*Preference Changes* S can be conveniently graphed as slopes between option sets {A,B} and {C,D}; negative and positive slopes indicate S > 0 and S < 0, respectively (Fig. 2). The strongest positive *Preference Changes* S were significant across sessions in all animals: S = 0.026 ± 0.013 (Monkey A), S = 0.095 ± 0.015 (Monkey B), S = 0.052 ± 0.010 (Monkey C) (mean ± Standard Error of the Mean, SEM; all *P* < 0.05; one-sample t-test) (Fig. 2*A*). The strongest negative S’s were also significant across sessions in all animals: S = -0.254 ± 0.020 (Monkey A), S = -0.167 ± 0.014 (Monkey B), S = -0.123 ± 0.014 (Monkey C) (Fig. 2*B*). The weakest absolute S’s differed only insignificantly from zero and thus failed to demonstrate IA violation (Fig. 2*C*). Fig. S1 shows the full pattern of *Preference Changes* in all CC tests.

In CR tests, *Preference Changes* are only necessary and not sufficient for IA violations; the test places gambles B and D at different distances from the respective parallel indifference lines (Fig. 1*D*), reflecting different expected utility differences for the two option sets that may result in graded *Preference Changes* and thus non-zero S values (but not outright *Preference Reversals*) but do not indicate IA violations. The strongest positive *Preference Changes* S were significant across sessions: S = 0.095 ± 0.0143 (Monkey A), S = 0.014 ± 0.015 (Monkey B), S = 0.078 ± 0.009 (Monkey C); all *P* < 0.05) (Fig. 3*A*), as were the strongest negative S’s: S = -0.183 ± 0.014 (Monkey A), S = -0.227 ± 0.019 (Monkey B), S = -0.086 ± 0.012 (Monkey C) (Fig. 3*B*). The smallest measured absolute S’s were insignificant (Fig. 3*C*). Fig. S2 shows the full pattern of *Preference Changes* in all CR tests.

To summarize, all monkeys showed significant *Preference Changes* in both CC and CR tests. Below we describe these results in more detail to identify possible factors contributing to the observed patterns of *Preference Changes*.

### Probability dependency of *Preference Change*s

Whereas previous human studies tested specific gambles, behavioral studies with monkeys can last several months during which large numbers of behavioral tests can be carried out. We have therefore been able to study choices of gambles over larger ranges of probabilities that fill wider areas of the Marschak-Machina triangle. This possibility allowed us to test whether the *Preference Changes* might depend on the probabilities of gamble outcomes, irrespective of particular preferences between the initial gambles A and B.

For the CC test, we varied the probability of the low outcome of gamble B (*Bp1*; i.e. the probability of receiving 0 ml in option B) and the probability of the high outcome (*Bp3*; i.e. the probability of receiving 0.5 ml in option B; thus, *Bp2* = 1 - *Bp3* - *Bp1*). In accordance with the definition of the CC test, we defined gambles C and D by adding a common probability k to options A and B (Fig. 1*C*). This corresponded to adding probability *Bp2* to the probabilities *p1* of gambles A and B (and thus reducing the original *p2* of gambles A and B). Therefore, we fully identified each CC test by the set of probabilities for gamble B (*Bp1, Bp*3), without the need to explicitly introduce the probability k. Significant IA *Preference Changes* occurred in both directions in different parts of the Marschak-Machina triangle (Fig. 4*A*; S > 0, purple dots; S < 0, cyan dots; black circles indicate *P* < 0.05, one-sample t-test against zero). As shown in the confusion matrices, LDA classifications correctly predicted 14 out of 18 tests (78%) in each of Monkey A and Monkey B, and 11 out of 18 tests (61%) in Monkey C, which exceeded random prediction (50%) and was no less than prediction with majority class (i.e. the majority type of the direction of *Preference Change* for each monkey; 72% for Monkey A, 50% for Monkey B and 61% for Monkey C) (Fig. 4*A* insets). These results suggested a systematic relationship between *Preference Changes* and reward probabilities in the CC test.

For the CR test, we varied the ratio *r* and the high-outcome probability in gamble B (*Bp3*; i.e. the probability of receiving 0.5 ml in option B; thus, *Bp1* = 1 - *Bp3*). We defined gambles C and D by multiplying the common ratio r with the probabilities of all non-zero outcomes of gambles A and B. Therefore, the two variables *Bp3* and *r* defined fully a particular CR test. Significant IA *Preference Changes* occurred in both directions in different parts of the parameter space (Fig. 4*B*; S 0, purple dots; S < 0, cyan dots; *P* < 0.05). The confusion matrices showed that LDA classifications correctly predicted 17 out of 18 tests (94%) in Monkey A, 15 out of 18 tests (83%) in Monkey B, and 15 out of 17 tests (88%) in Monkey C, all of which exceeded random prediction and was no less than prediction with majority class (94% for Monkey A, 67% for Monkey B, 82% for Monkey C) (Fig. 4*B* insets). Hence, similar to the CC test, the *Preference Changes* in the CR test depended on gamble probabilities.

The systematic nature of the observed *Preference Changes* in both the CC tests and the CR test encouraged us to model the observed changes mathematically using economic theory.

Therefore, we next fitted our data using different economic choice models and tested whether these models might explain the observed violations.

### Economic models characterizing *Preference Changes*

We compared different economic choice models in their ability to explain the observed pattern of Preference Changes. We fitted choice data to stochastic implementations of basic constructs of three economic theories: objective Expected Value (EV), Expected Utility Theory (EUT) and Cumulative Prospect Theory (CPT).

We defined a standard discrete choice softmax function (McFadden, 2001) to describe stochastic preferences as the probability of choosing one option over another, in repeated trials (Eq. 7). This function calculates the probability of choosing the first of two options from the value difference between the two options and includes a noise term that accounts for variability in choices. The difference between the choice models consisted of different value computations: in the EV model, each option’s value corresponded to its objective Expected Value (EV; Eqs. 8 and 9); in the EUT model, value was defined as Expected Utility using a utility function (EU; Eqs. 10 and 11); in the CPT model, value was defined as *Prospect Value* and resulted from a utility function (*u*) and a probability weighting function (*w*), combined in a cumulative form (Eqs. 12-14). The utility function was a power function (one free parameter), normalized to the highest magnitude (Eq. 15). The probability weighting function was a two-parameter *Prelec* function (Eq. 16). These parametric functions have been shown to maximize the information extraction from participant data (Stott, 2006). Finally, we used a maximum likelihood estimation procedure to identify the model parameters that best represented the behaviorally measured probability of choice: we estimated the parameters that maximized the standard log-likelihood function (Eq. 17).

We used the Akaike Information Criterion (AIC) for an initial comparison of the accuracy of each model, based on the maximum likelihood function (lower AIC values indicate better fit). The AICs of the EV model were 204.4 ± 11.4 for Monkey A, 144.8 ± 4.8 for Monkey B, and 216.8 ± 8.0 for Monkey C (mean ± standard error of the mean, SEM). The AICs of the EUT model across sessions were 149.5 ± 10.8 for Monkey A, 118.7 ± 4.2 for Monkey B, and 115.6 ± 4.2 for Monkey C. The AICs of the CPT model were 140.5 ± 10.1 for Monkey A, 114.4 ± 4.1 for Monkey B, and 110.4 ± 4.3 for Monkey C. The differences between the three AIC values were significant in each animal (*P* = 6.08·10^−05^ for Monkey A, *P* = 1.50·10^−06^ for Monkey B, and *P* = 4.67·10^−33^ for Monkey C; one-way ANOVA). Pairwise post-hoc comparison showed significant differences between the EUT and EV models (*P* = 2.98·10^−16^ for Monkey A, *P* = 8.47·10^−20^ for Monkey B, and *P* = 5.55·10^− 30^ for Monkey C; paired t-test) and between the CPT and EUT models (*P* = 8.41·10^−10^ for Monkey A, *P* = 2.02·10^−10^ for Monkey B, and *P* = 8.58·10^−11^ for Monkey C). Thus, the CPT model showed the lowest AIC values in all three monkeys and thus explained our choice data best. We therefore used the CPT model for our further analyses.

According to the CPT model, Monkey A had basically a linear utility function (U(m); estimated parameter: *ρ* = 1.01) and a probability weighting function with an inverse-S shape (W(p); estimated parameters: *α* = 1.86; *β* = 0.42), whereas Monkeys B and C had convex utility functions (*ρ* = 1.43 and *ρ* = 1.63, respectively) and inverted-S-shaped probability weighting functions (*α* = 1.31; *β* = 1.13 and *α* = 1.37; *β* = 0.767, respectively) (Fig. 5*A*, insets).

To better understand and visualize how the CPT model might explain the IA violations, we computed the indifference curves (ICs) in the Marschak-Machina triangle (Fig. 5*A* left), based on the utility and probability weighting functions estimated from the best-fitting CPT model (Fig. 5*A*, insets). According to the EV and EUT models, the ICs in the Marschak-Machina triangle should be linear and parallel to each other, while CPT produces non-linear and non-parallel ICs. The indifference map (i.e. the full set of ICs) computed from the best fitting CPT model in each animal showed monkey-specific patterns of non-linear ICs, which reflected the subjective value of choice options (Fig. 5*A*, colored lines). We considered the “fanning” direction of the ICs to further characterizes IA violations (Machina, 1982); “fanning-out” (higher ICs more horizontal than lower ones) characterizes Allais-type violations, and “fanning-in” characterizes reverse Allais-type violations. We observed a predominantly fanning-in pattern, although areas of fanning-out existed within the triangle. This pattern reflected the mostly negative values of the measured *Preference Changes*, supporting the idea that IA violations reflected a non-linear distribution of subjective values within the Marschak-Machina triangle, which is incompatible with EUT.

To examine how well CPT explained the observed *Preference Changes*, we calculated the S values that were predicted by the model for each CC and each CR test. On a session-by-session basis, we estimated the choice probability from outcome probability and magnitude according to Eq. 7 together with Eqs. 12 - 14. The estimated choice probability was then used to calculate each S using Eq. 6. When comparing the measured and predicted S values for each session, we found significant Pearson correlation coefficients in all monkeys in the CC test (Monkey A: *ρ* = 0.46, *P* = 2.1·10^−53^; Monkey B: *ρ* = 0.21, *P* = 8.9·10^−9^; Monkey C: *ρ* = 0.29, *P* = 4.4·10^−49^) as well as in the CR test (Monkey A: *ρ* = 0.74, *P* = 1.1·10^−188^; Monkey B: *ρ* = 0.60, *P*= 2.4·10^−72^; Monkey C: *ρ* = 0.64, *P* = 2.6·10^−287^) (Fig. 5*B*). Thus, the CPT model was compatible with the observed pattern of violations.

We tested the robustness of the CPT model’s IC estimation with out-of-sample tests. The animal chose between a fixed option, plotted on one of the axes of the Marschak-Machina triangle, and a varying two- or three-outcome gamble (see Methods). In each session, indifferent points (IPs) were estimated by fitting a softmax function to the measured animal’s choices. If the modeled indifference map reflected the true subjective evaluation pattern, the modeled IPs should be close to the measured ICs. A graphical comparison between the IPs and the ICs in Fig. 6*A* predicted by the CPT model demonstrated good correspondence between the out-of-sample IPs (colored points) and the modeled ICs (lines with same color as IPs). When quantifying the distance between the measured out-of-sample IPs and the ICs predicted by the EV, EUT or CPT models, we found significant residuals in all three models (*P* < 0.01 against the ICs; one-sample t-test). The residuals were significantly different across the models in all three monkeys, as revealed by one-way ANOVA tests (*P* = 3.10·10^−86^ for Monkey A, *P* = 2.66·10^−33^ for Monkey B, and *P* = 5·10^−324^ for Monkey C); post-hoc paired t-tests demonstrated smaller residuals for the CPT model compared to the EV and EUT models (Fig. 6*B*), except for data from Monkey B resulting in a non-significant difference between EUT and CPT model residuals (EUT vs EV: *P* = 5.24·10^−26^ for Monkey A, *P* = 2.84·10^−32^ for Monkey B, and *P* = 1.35·10^−134^ for Monkey C; CPT vs EUT: *P* = 1.28·10^−08^ for Monkey A, *P* = 0.734 for Monkey B, and *P* = 9.26·10^−22^ for Monkey C). Thus, the CPT model captured the out-of-sample IPs more accurately than the other models.

**Fig. 6.**
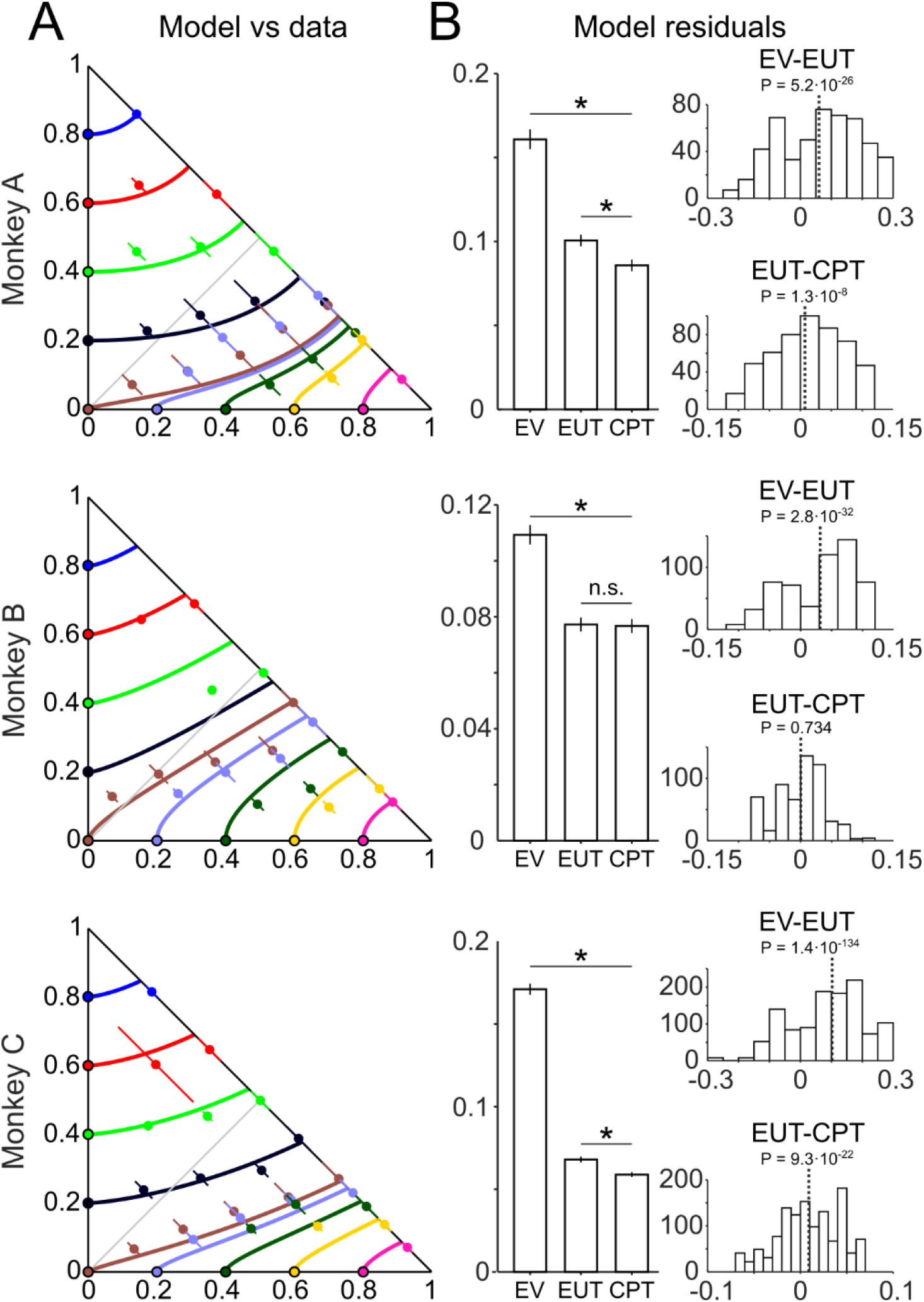
Out-of-sample tests on indifference curves (ICs) modeled by Cumulative Prospect Theory (CPT). (A) Close relationship between measured out-of-sample indifference points (IP, colored dots) and ICs modeled by CPT (colored lines) in common ratio and common consequence tests. Colored dots show mean IPs across all sessions, corresponding to the same-colored ICs; lines show Standard Deviations (SD) of IPs across all sessions. (B) Left: bar charts of means and Standard Errors of the Mean (SEM) of model residuals (distances between model ICs and the measured out-of-sample IPs). Asterisks: *P* < 0.05 in post-hoc paired t-test (n.s.: not significant, *P* > 0.05). Right: residual differences from different models (means from individual sessions; top: Expected Value (EV) minus Expected Utility Theory (EUT); bottom: EUT minus CPT. Dotted line: mean value; *P*: post-hoc paired t-test p-value. Smaller residuals for CPT than EUT or EV indicate that CPT better captured the pattern of the measured out-of-sample IPs.

Although in their original deterministic formulation the EV and EUT models would not theoretically produce all-or-none *Preference Reversals*, their stochastic versions using the softmax choice function could in principle result in graded *Preference Changes*, in particular in the CR test (see Methods below Eq. 6). We thus investigated in more detail how much the EUT and EV models would explain our data (Figs. S3, S4). Our analysis showed that EV and EUT models failed to explain violations in the CC test, always predicting null *Preference Changes* (S = 0) (Figs. S3*B*, inset). On the other hand, both models predicted the violation pattern in the CR test to some degree (Figs. S3*B*, S4*B*), consistent with previous studies employing stochastic versions of the EUT model (P. R. Blavatskyy, 2007). However, the Pearson correlation coefficients of the EV and EUT models had worse prediction power (smaller correlation coefficients) compared to the CPT model (EV: *ρ* = 0.26, *P* = 3.1·10^−18^ for Monkey A, *ρ* = 0.51, *P* = 9.3·10^−50^ for Monkey B, and *ρ* = 0.22, and *P* = 1.7·10^−28^ for Monkey C; and EUT: *ρ* = 0.55, *P* = 4.4·10^−86^ for Monkey A, *ρ* = 0.59, *P* = 2.1·10^−69^ for Monkey B, and *ρ* = 0.52, *P* = 8.6·10^−172^ for Monkey C) (Figs. S3 and S4).

As a further control, we explicitly tested the hypothesis of ICs being linear and parallel, as implied by EUT (Fig. S5). Because previous human studies usually performed tests on only a few gambles, as plotted in the Marschak-Machina triangle (Fig. 7*A*; Blavatskyy et al., 2015), this method has never been used to investigate EUT. In the current study, we tested separately the linearity and parallelism of the ICs. To test for parallelism, we assumed linear ICs and used a least-squares model to estimate the slopes of the ICs and compare them (Kruskal-Wallis one-way ANOVA; Fig. S5*A, B*). The linearity of the ICs was tested through the residuals of indifferent points in each IC that were estimated with linear least squares using out-of-sample IPs (one-sample t-test against 0; Fig. S5*C*). We found significant non-linearity (p<0.001) and non-parallelism (p<0.05) for some ICs, suggesting that EUT was not able to capture the subjective values for varying probabilities. This result demonstrates systematic violations in EUT, which was consistent with our AIC and residual analyses. The pattern of generally increasing ICs slopes (Fig. S5*B*) also served as confirmation for a mostly fanning-in direction of the indifference map, which explained the observed pattern of IA violations with mostly negative *Preference Change* values.

**Fig. 7.**
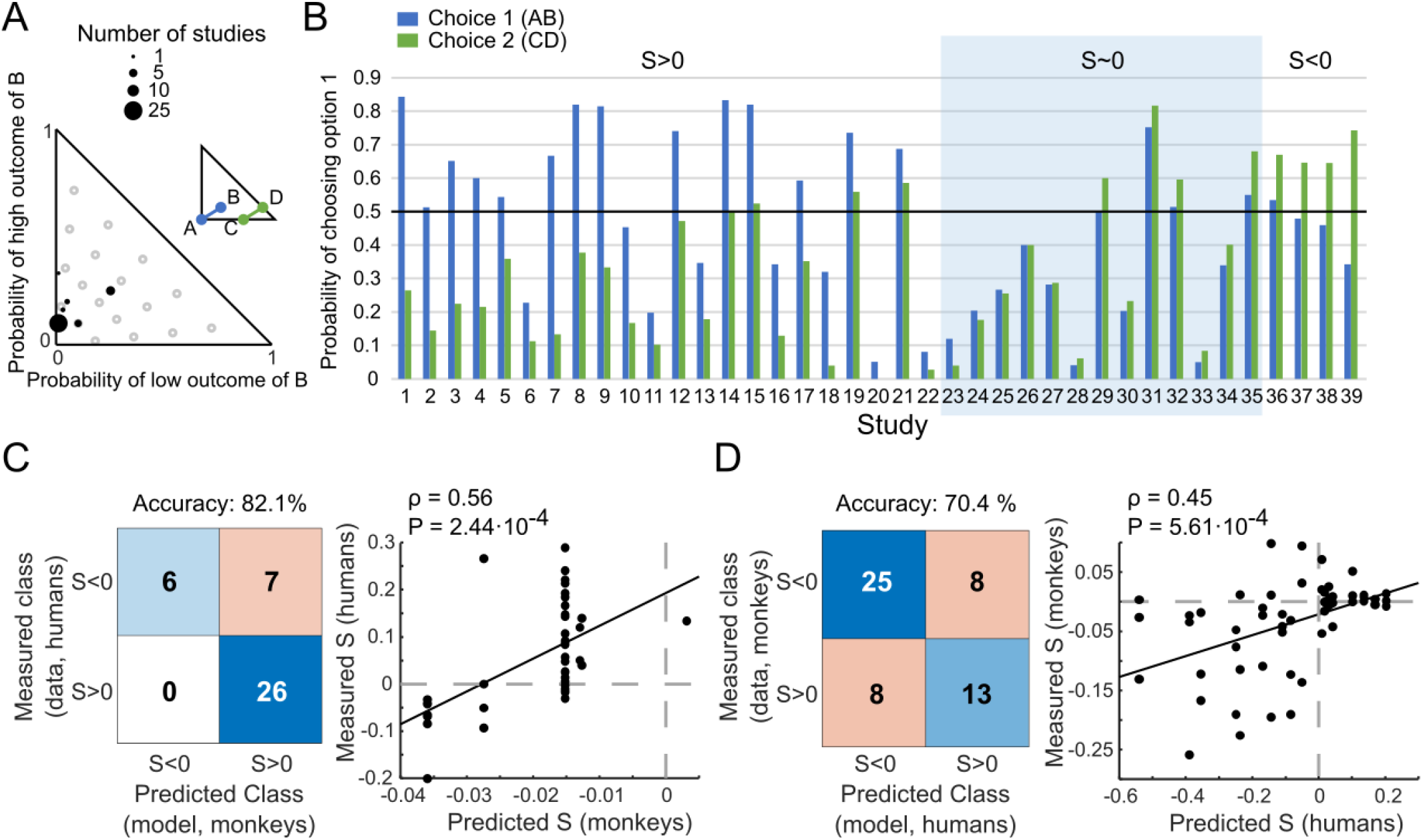
Correspondence of *Preference Changes* S between monkeys and humans. (A) Gamble positions in the Marschak-Machina triangle of 39 independent human common consequence studies (P. Blavatskyy et al., 2015). The x-axis represents the probability of getting the low outcome in option B and the y-axis represents the probability of getting the high outcome in option B (see option B in Fig. 1*D*). The diagram illustrates the location of the reward probability tested in the human studies (black dots). Gray circles correspond to the CC tests we performed on monkeys in the current study. (B) Results from the 39 human studies. Blue and green bars refer to option sets 1 {A, B} and 2 {C, D}, respectively (see Fig. 1*B*). The y-axis indicates the probability of choosing option 1 (A or C). (C) Correspondence between measured human S’s (39 studies; *B*) and predicted monkey S’s. Left: confusion matrix of classes of human S’s and classes of monkey S’s predicted from actually used monkey gambles by Linear Discriminant Analysis (LDA). The LDA prediction of monkey S’s allowed comparison of same gambles between the two species. Right: Pearson correlation between measured human S’s and monkey S’s predicted by regression (Eq. 18, 19), using the same gambles. (D) Correspondence between measured monkey S’s and predicted human S’s. This inverse control test relative to the test shown in *C* was based on the actual gambles used in monkeys and employed predictions of human S’s for these gambles via different LDA and regression (Eqs. 20, 21; see Methods).

Taken together, these results showed that the CPT-based economic choice model predicted the IA violations in both CC and CR tests, outperforming both the EV and the EUT models. These findings suggest that the observed violation pattern might arise from the subjective, non-linear weighting of reward probabilities, in line with the explanation provided by CPT.

### Comparison of *Preference Changes* with humans

To explore the possibility of common economic decision mechanisms between evolutionary close species, we compared the observed IA violations in monkeys with those found in humans. We considered results from 39 human studies investigating the CC effect (P. Blavatskyy et al., 2015). Many of these studies repeated the Allais test; others defined different tests which, when represented in the Marschak-Machina triangle (as *p1* and *p3* of gamble B), were mostly concentrated in the lower left area (Fig. 7*A*). The human studies reported significant *Preference Changes* characterized by S > 0 or S < 0, as well as insignificant changes (S ∼ 0) (Fig. 7*B*).

We used two methods to compare our monkey data with the published human data, a confusion matrix and a Pearson correlation. The gambles used in the human studies differed from each other and from those used in monkeys. To nevertheless allow accurate comparisons, we predicted the *Preference Changes* S for the untested gambles with an LDA classifier for the confusion matrix, and with two multiple linear regressions for the Pearson correlation (see Methods; Eqs. 18 - 21).

For the missing gambles, we first used the LDA and the regressions (Eqs. 18 and 19) to predict the S’s in our monkeys for the gambles that had been used in humans. Then we compared the actually measured human S’s with the predicted monkey S’s. The confusion matrix showed that the actual measured S’s in humans corresponded successfully to the LDA-predicted S’s in monkeys with 82% accuracy (Fig. 7*C* left), which exceeded random (50%) and majority class (i.e. the majority type of the direction of S’s across all human studies, 67%). The Pearson correlation between human S’s that had been measured and monkey S’s that had been predicted by regression (Eqs. 18 and 19) was significant (*ρ* = 0.56, *P* = 2.44·10^−4^) (Fig. 7*C* right).

To test the robustness of these comparisons, we reversed the direction of predicting S’s for untested gambles: using LDA and regressions (Eqs. 20 and 21), we predicted the S’s in humans for the gambles we had used in monkeys. The confusion matrix showed that the measured monkey S’s corresponded to the LDA-predicted human S’s with 70% accuracy, which exceeded random (50%) and majority class (61%) (Fig. 7*D* left). The Pearson correlation between the measured monkey S’s and the regression-predicted human S’s (Eqs. 20 and 21) was significant (*ρ* = 0.45, *P* = 5.61·10^−4^; Fig. 7*D* right).

Thus, while we saw less *Preference Reversals* than are generally reported in humans, the *Preference Changes* in our monkeys corresponded well to those in humans. The result suggests shared decision mechanisms across primate species and encourages neurophysiological investigations in monkeys of neuronal signals and circuits that may underlie these common choice mechanisms.

## Discussion

We studied stochastic choices in rhesus monkeys in the two most widely used tests of the IA, common consequence and common ratio, which provide stringent assessment for utility maximization. All three tested monkeys showed consistently few outright *Preference Reversals* between the initial and the altered option sets, possibly due to the animals’ extended laboratory experience with weekdaily tests; however, the animals showed substantial graded *Preference Changes* (Figs. 2 and 3; Tables 1 and 2) that depended on gamble probabilities and were largely explained by nonlinear probability weighting (Figs. 4 and 5). According to AIC and out-of-sample analyses, a CPT model with probability weighting explained the choices better than EUT and EV models without probability weighting (Fig. 6). Classification and regression analyses demonstrated similarities between our monkeys’ choices and reported human choices (P. Blavatskyy et al., 2015, 2022) (Fig. 7). Together, these results indicate systematic *Preference Changes* in IA tests in monkeys that can be explained by probability weighting of CPT.

While human studies played a crucial role in identifying and explaining IA violations, in particular non-linear probability weighting (P. Blavatskyy et al., 2022; P. R. Blavatskyy, 2007; Camerer & Ho, 1994; Conlisk, 1989; Harman & Gonzalez, 2015; Quiggin, 1982; Ruggeri et al., 2020; Schneider & Day, 2016; Tversky & Kahneman, 1992), the studies were restricted by a number of species-specific factors, including limited trial numbers, limited test variations, limited test repetitions, insufficient learning, behavioral errors and, of course, language and cultural influences. To compensate for limits of trial numbers, some human studies combined data from multiple participants; however, the validity of such tests depends on the subjectivity of individual utility functions and on cultural differences (Loubergé & Outreville, 2011; Ruggeri et al., 2020). Thus, more comprehensive assessments of IA compliance would benefit from wider test variations with more trials than are feasible in humans. This is where monkeys come in. Working with monkeys not only avoids cultural biases but also allows large variations of test conditions during thousands and tens of thousands of trials during weekdaily tests over weeks and months. With such large trial numbers, errors are minimized and learning would be completed and thus play no uncontrolled role. The resulting consistent performance allowed us to investigate the robustness of economic models that confirm the dominant role of probability weighting in common consequence and common ratio IA tests.

This study found fewer outright *Preference Reversals* (8%) than those seen in human studies; we saw primarily graded *Preference Changes*. The limited violations in the IA tests resemble the compliance of the Independence of Irrelevant Alternatives of two-component bundles (Pastor-Bernier et al., 2017). Good compliance in the two tests may be due to the high task familiarity of the animals tested in thousands of trials. To assure well-controlled test conditions, our monkeys performed in our specific primate testing laboratory away from their living area in the animal house. For ethical reasons, such laboratory tests are limited to a few animals. However, in this highly standardized test situation the different animals performed very consistently and similarly to each other. In support of this notion, further four monkeys in two separate studies in our laboratory showed consistent risk attitude that was compatible with S-shaped, convex-linear-concave utility functions with increasing juice volumes (Genest et al., 2016; Stauffer et al., 2015).

Thus, while ethical considerations, general welfare and individual comfort are essential for obtaining reliable results from cooperative animals, the presented research on monkeys adds important data to the notion of assessing utility maximization with the IA axiom that had so far been tested in humans and, in select cases, in rodents (Battalio, Kagel, & MacDonald, 1985; Kagel et al., 1990).

Probability weighting is a particularly interesting and important explanation for IA violations. Although reward probability can influence the type and level of IA violations, most human IA violation tests used only limited levels of probability. We tested many probability levels across the Marschak-Machina triangle and substantiated probability weighting as major mechanism underlying *Preference Changes* in both the common consequence and the common ratio tests. We did not make any hypothesis about the existence of probability weighting but instead demonstrated empirically that probability weighting explains *Preference Changes*. Specifically, our leave-one-out classification with LDA demonstrate probability as key factor underlying IA *Preference Changes* in both common consequence and common ratio tests (Fig. 4), and the CPT probability weighting function fitted to the measured behavioral choices successfully predicted *Preference Changes* (Fig. 5). Thus, our study provides a detailed and robust account of the role of probability weighting in IA tests. In addition to probability weighting, it has been proposed that the salience of the visual cues for reward probability information (i.e., the length of the stimulus bar in our study) could contribute to choice biases (Spitmaan, Chu, & Soltani, 2019), suggesting a future direction for investigating its role in IA violations.

Past studies have reported different shapes of the probability weighting function. Humans show anti-S and S-shape probability weighting with instructed and experienced probabilities, respectively (Cavagnaro, Pitt, Gonzalez, & Myung, 2013; Farashahi, Azab, Hayden, & Soltani, 2018; Gonzalez & Wu, 1999). Monkeys show anti-S probability-weighting with pseudorandomly varying probabilities (Stauffer et al., 2015) and S-shape weighting in trial blocks (Ferrari-Toniolo et al., 2019). When presenting a larger set of magnitudes and probabilities, and allowing for more complex shapes of the utility function, our monkeys’ choices were best explained by a mostly concave probability weighting function (Ferrari-Toniolo et al., 2021). Humans show a similar concave probability weighting function (P. Blavatskyy, 2013). Our current results confirm concave probability weighting with a larger set of finely varying probabilities in three-outcome gambles that allowed us to uniformly sample the whole probability space (Fig. 5*A*). Our results highlight a series of possible factors contributing to the estimated shape of the probability weighting function: the choice of the functional form for utility and probability weighting, the range and resolution of the tested magnitudes and probabilities, and the complexity and representation of choice options (especially two- and three-outcome gambles). Further investigations are required to better isolate the factors influencing the shape of the probability weighting function, including task particulars and elicitation procedures.

Past studies graphed IA violations via the fanning-in and fanning-out directions of ICs in the Marschak-Machina triangle (Machina, 1982). Non-linear ICs, compatible with nonlinear probability weighting, produce different local fanning directions in different areas of the triangle (Fig. 5*A*) (Camerer & Ho, 1994; Kontek, Kontek, & Krzysztof, 2018; G. Wu & Gonzalez, 1998).

Furthermore, different stochastic versions of EUT (P. R. Blavatskyy, 2007) and, more in general, different contributions of noise to the value computation mechanism (Bhatia & Loomes, 2017; Hey & Orme, 1994; Woodford, 2012) might explain IA violations without nonlinear probability weighting. These considerations highlight the complexity in the relation between the shape of the probability weighting function, the pattern of indifference curves and the experimentally revealed types of IA violations. Further theoretical work and model simulations, which are outside of the scope of the current work, should help to elucidate these relations.

Human tests of the IA describe *Preference Changes* characterized by S > 0 or S < 0 (Allais, 1953; P. Blavatskyy et al., 2022; Starmer, 2000). Because of these violations, many economic theories have been developed to explain economic choices under risk, including Rank-Dependent Utility (Quiggin, 1982), Cumulative Prospect Theory (Tversky & Kahneman, 1992), and Target-

Adjusted Utility (Schneider & Day, 2016). Consistent with human choices, we found that our monkeys’ choices show both types of violations in the two most studied tests (common consequence and common ratio). Interestingly, although humans and monkeys may show different probability weighting functions, the *Preference Changes* S seen in the repeated, stochastic choices of our monkeys correspond with 70% - 82% accuracy to the S’s computed from choices averaged across human participants (Fig. 7*C, D*). This result is not only interesting for general inter-species comparisons but indicates that violations in primates are similar and robust despite methodological differences, such as trial numbers and averaging within subjects as opposed to across subjects.

The IA is arguably the most constrained and direct test that defines Expected Utility, and its maximization, on a numeric, cardinal scale. With these properties, the IA provides for a stringent test framework for investigating brain mechanisms of economic choice. So far, human fMRI studies demonstrate subjective value coding in reward-related brain regions, including the ventral striatum, midbrain, amygdala, and orbitofrontal and ventromedial prefrontal cortex (Gelskov, Henningsson, Madsen, Siebner, & Ramsøy, 2015; Hsu, Krajbich, Zhao, & Camerer, 2009; Seak, Volkmann, Pastor-Bernier, Grabenhorst, & Schultz, 2021; S.-W. Wu, Delgado, & Maloney, 2011).

Neurophysiological studies in monkeys demonstrate the coding of subjective value in midbrain dopamine neurons and orbitofrontal cortex (Kobayashi & Schultz, 2008; Lak, Stauffer, & Schultz, 2014; Padoa-Schioppa & Assad, 2006; Stauffer et al., 2014; Tremblay & Schultz, 1999) and formal utility coding in dopamine neurons (Stauffer et al., 2014). Further, neurons in monkey orbitofrontal cortex carry single-dimensional utility signals for two-dimensional choice options designed according to Revealed Preference Theory (Pastor-Bernier, Stasiak, & Schultz, 2019). However, despite attempts of economic decision theories to explain IA violations (such as prospect theory), the neuronal mechanisms underlying IA violations are unknown. To address the issue, an animal model would be desirable that demonstrates IA violations similar to those seen and analyzed in humans. Our own studies showed that monkeys’ choices follow indifference curves of Revealed Preference Theory, satisfy first-, second- and third-order stochastic dominance, demonstrate probability weighting, and comply with the first three EUT axioms (completeness, transitivity, continuity) (Ferrari-Toniolo et al., 2021, 2019; Genest et al., 2016; Pastor-Bernier et al., 2017; Stauffer et al., 2015), all of which suggests compliance with fundamental concepts of economic choice. The current study describes compliance and violation of the fourth EUT axiom, IA, which is the most demanding and investigated EUT axiom in humans. Tests in rodents have revealed globally similar IA violations as in humans (Battalio et al., 1985; Kagel et al., 1990), but the results have so far not been used for neurophysiological investigations in this species. As the performance of our monkeys in IA tests is also consistent with that in humans, researchers may want to use neurophysiology in animals to understand neuronal choice mechanisms in humans.

Although our study provides systematic and stochastic data on IA violations, there are a few incompletely addressed directions that can be investigated in the future. For example, further research may test whether reward magnitude can influence IA violations, as it does in humans (P. Blavatskyy et al., 2022). Further, in the absence of own human data, we can only relate our results to those from human experiments that did not necessarily have the exact same design. Some of the observed differences between human’s and monkey’s IA violations could be due to the unequal sampling of the probability space across species, with human studies usually focusing on a specific region of the Marschak-Machina triangle. Our analyses revealed a difference in the magnitude of the S values between species, together with a minority of incompatible predictions for the *Preference Change* direction (Fig. 7*C, D*; confusion matrix and correlation plots). These differences might reflect the fact that our evaluations depended on the indirect comparison between measured and predicted Colinsk’ S values across species. To more directly compare IA violations between humans and monkeys (in which neurophysiological studies are more feasible), future studies might adapt our experimental design to that used in humans. We also observed some differences in effect size between positive and negative S values (Fig. 2), which suggest future investigations of distinct behavioral strategies leading to violations in different directions. Additionally, future experiments could investigate different classes of economic models that might capture more reliably the pattern of IA violations when allowing for the stochasticity of choice (P. R. Blavatskyy & Pogrebna, 2010; Loomes & Pogrebna, 2014). Our tests might support further development of decision theory and computer algorithms, for example by using our data for advancing model-free and model-based reinforcement learning theory into the domain of economic choice research (Daw, Gershman, Seymour, Dayan, & Dolan, 2011; Miranda, Malalasekera, Behrens, Dayan, & Kennerley, 2020). It would be interesting to see how subjective values are updated after win or loss trials in IA violated gambles (model-free: based only on stimuli; model-based: update the whole probability and utility model; or a combination of both). Neurophysiology research on value updating by reinforcement could benefit from the developed experimental designs. Thus, because of its multidisciplinary nature, our current behavioral study may provide the basis for further investigations of behavioral and neuronal mechanism of economic decision-making under risk.

## Acknowledgements

We thank Ms. Christina Thompson, Mr. Aled David and Dr. Henri Bertrand for animal and technical support. The Wellcome Trust (WT 095495, WT 204811), the European Research Council (ERC; 293549) and the US National Institutes of Mental Health Conte Center at Caltech (NIMH; P50MH094258) supported this work.

## Supplementary Figures for

**Supplementary Fig. 1.**
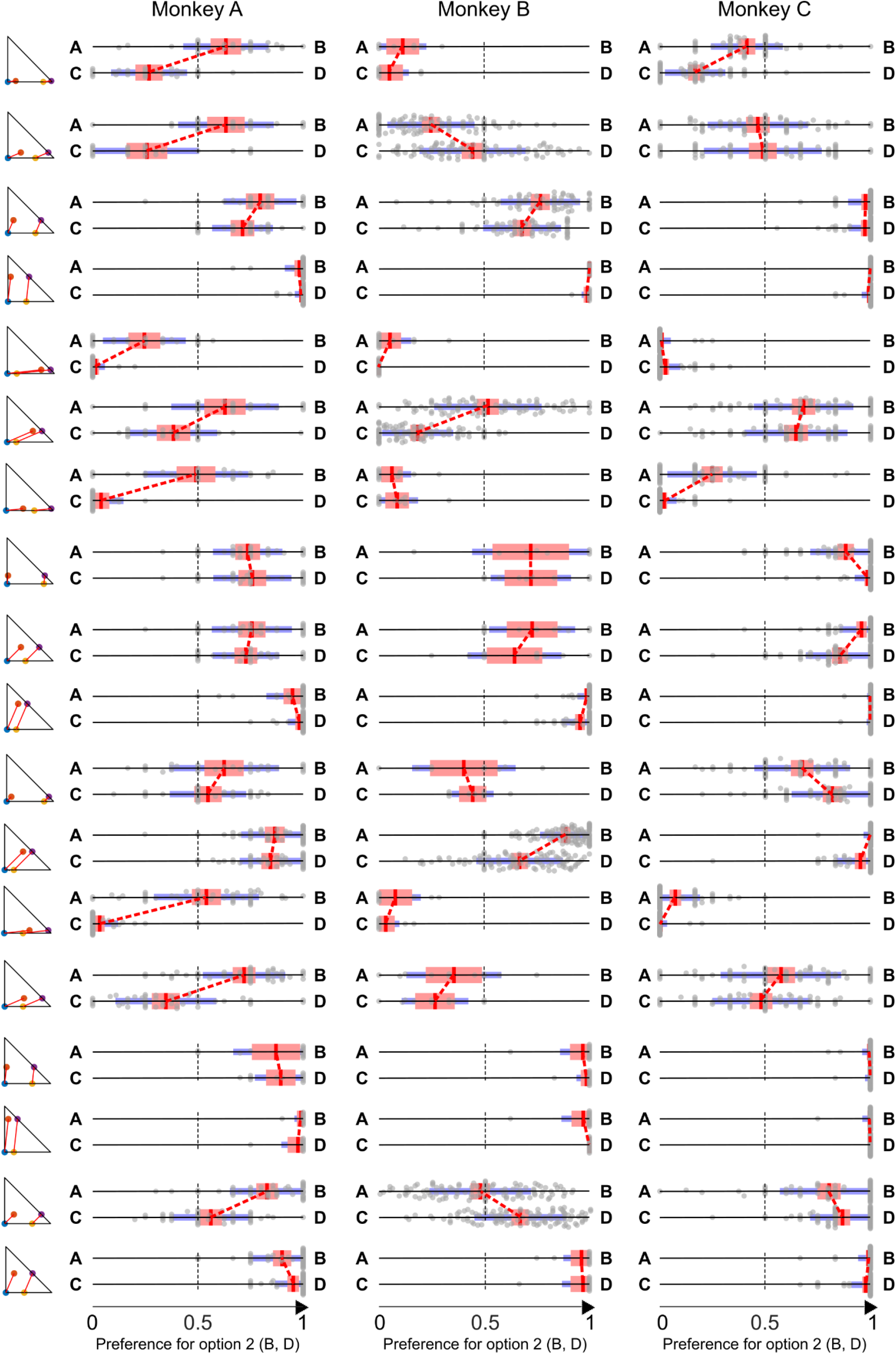
*Preference Changes* S for all common consequence tests. For conventions, see Fig. 2.

**Supplementary Fig. 2.**
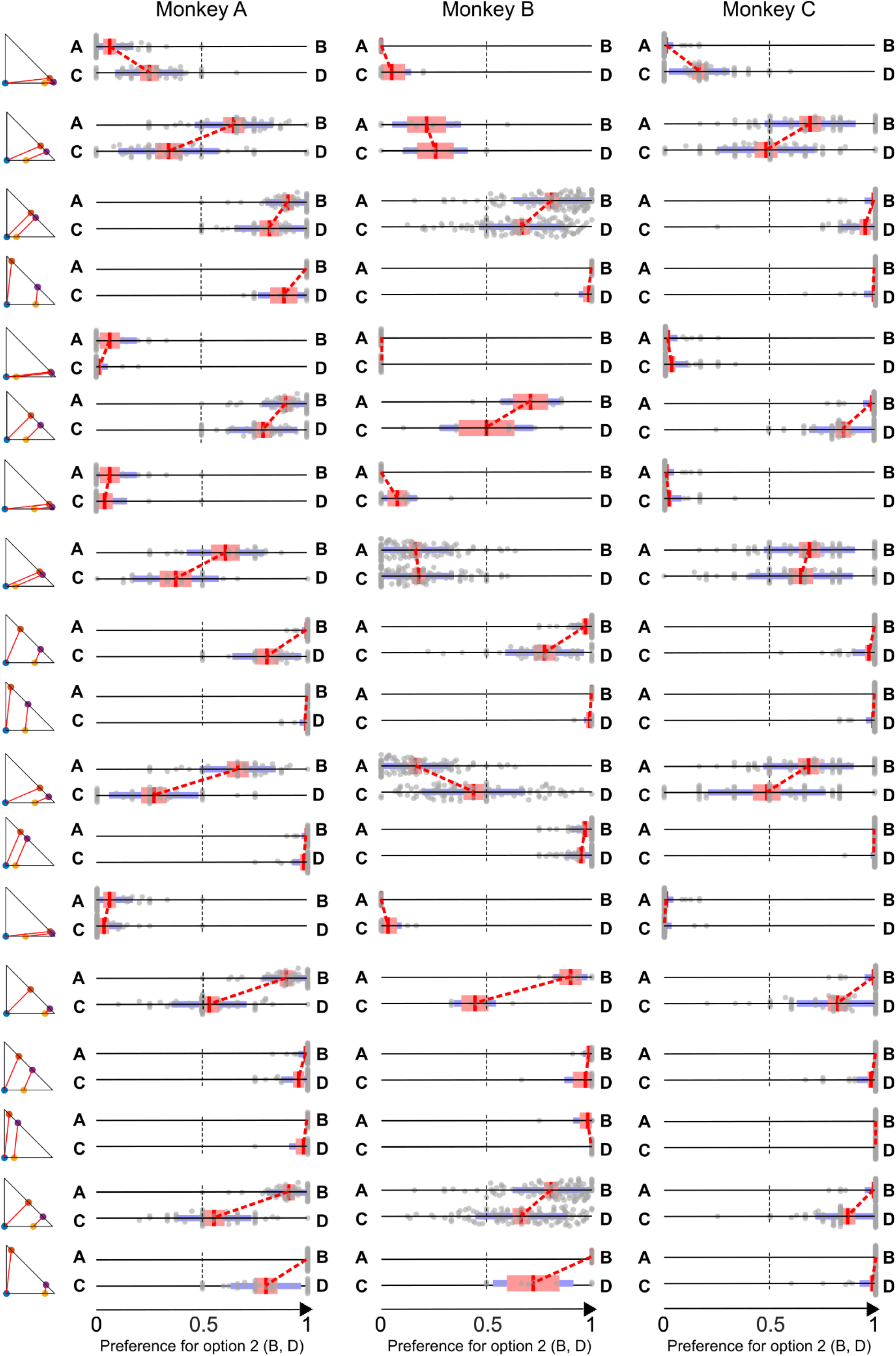
*Preference Changes* S for all common ratio tests. For conventions, see Fig. 2.

**Supplementary Fig. 3.**
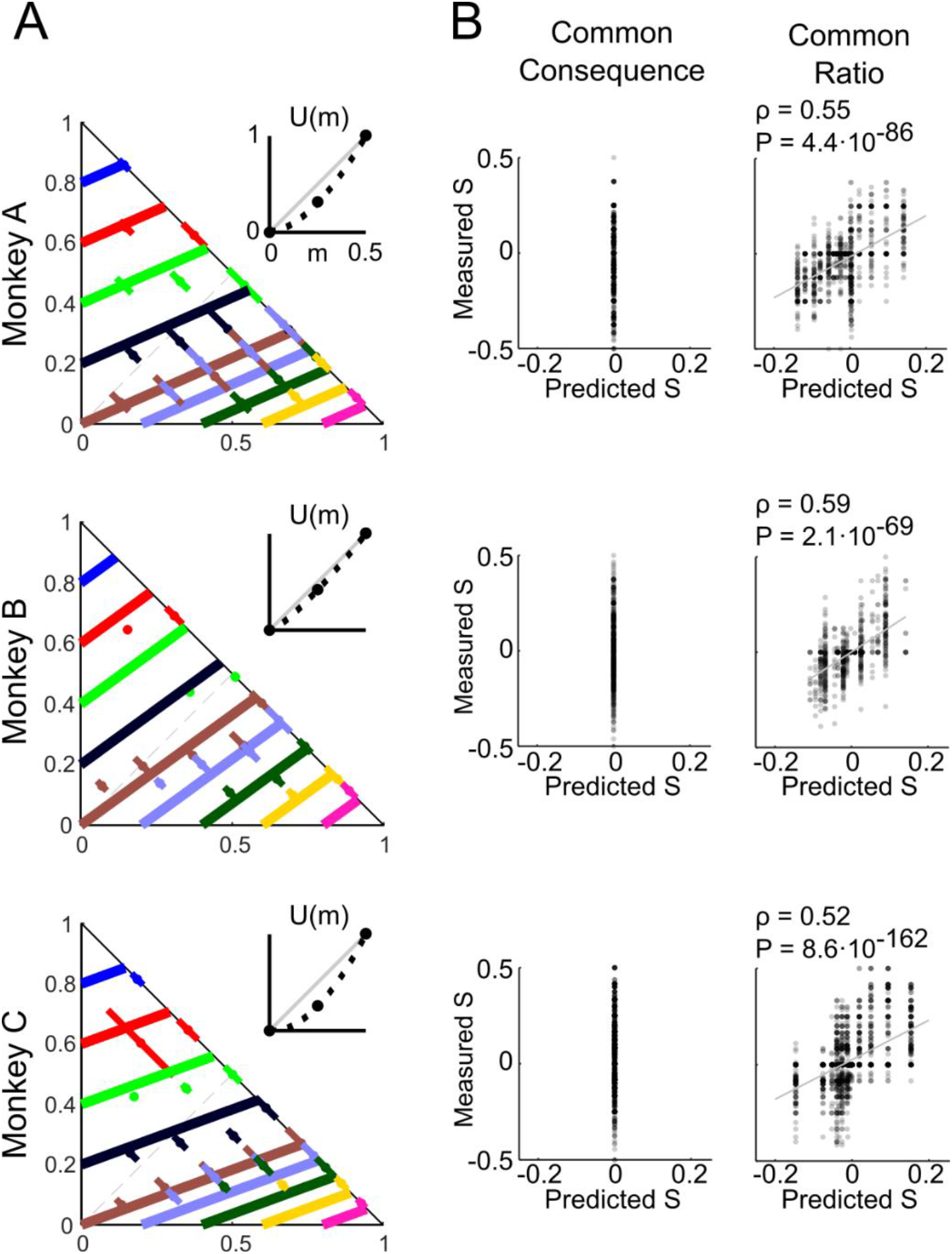
Expected Utility Theory (EUT) modeling failed to explain the common consequence (CC) test but can predict the Preference Changes of the common ratio (CR) test to some degree. For conventions, see Fig. 6.

**Supplementary Fig. 4.**
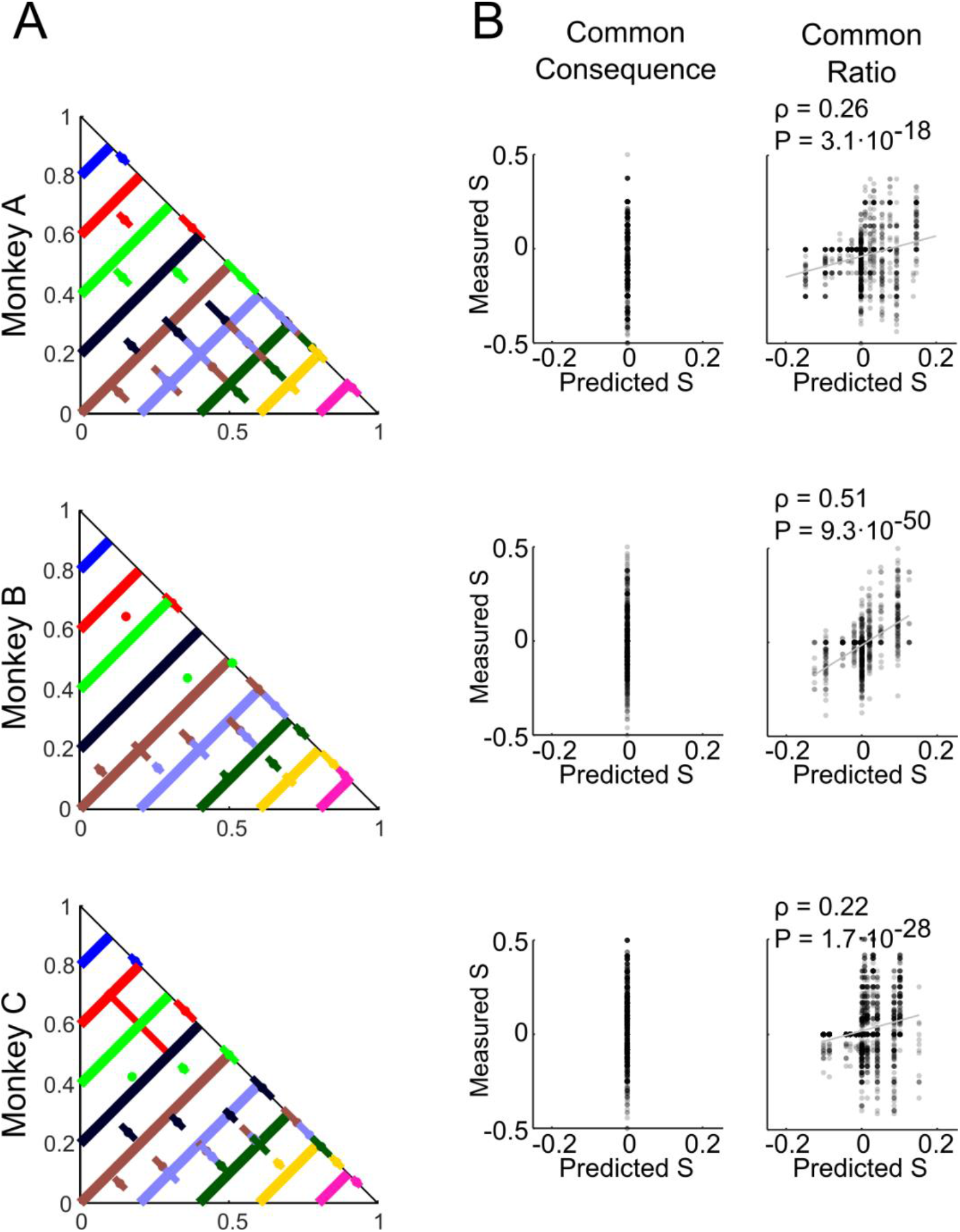
Expected Value (EV) modeling failed to explain the common consequence (CC) test but can predict the Preference Changes of the common ratio (CR) to some degree. For conventions, see Fig. 6.

**Supplementary Fig. 5.**
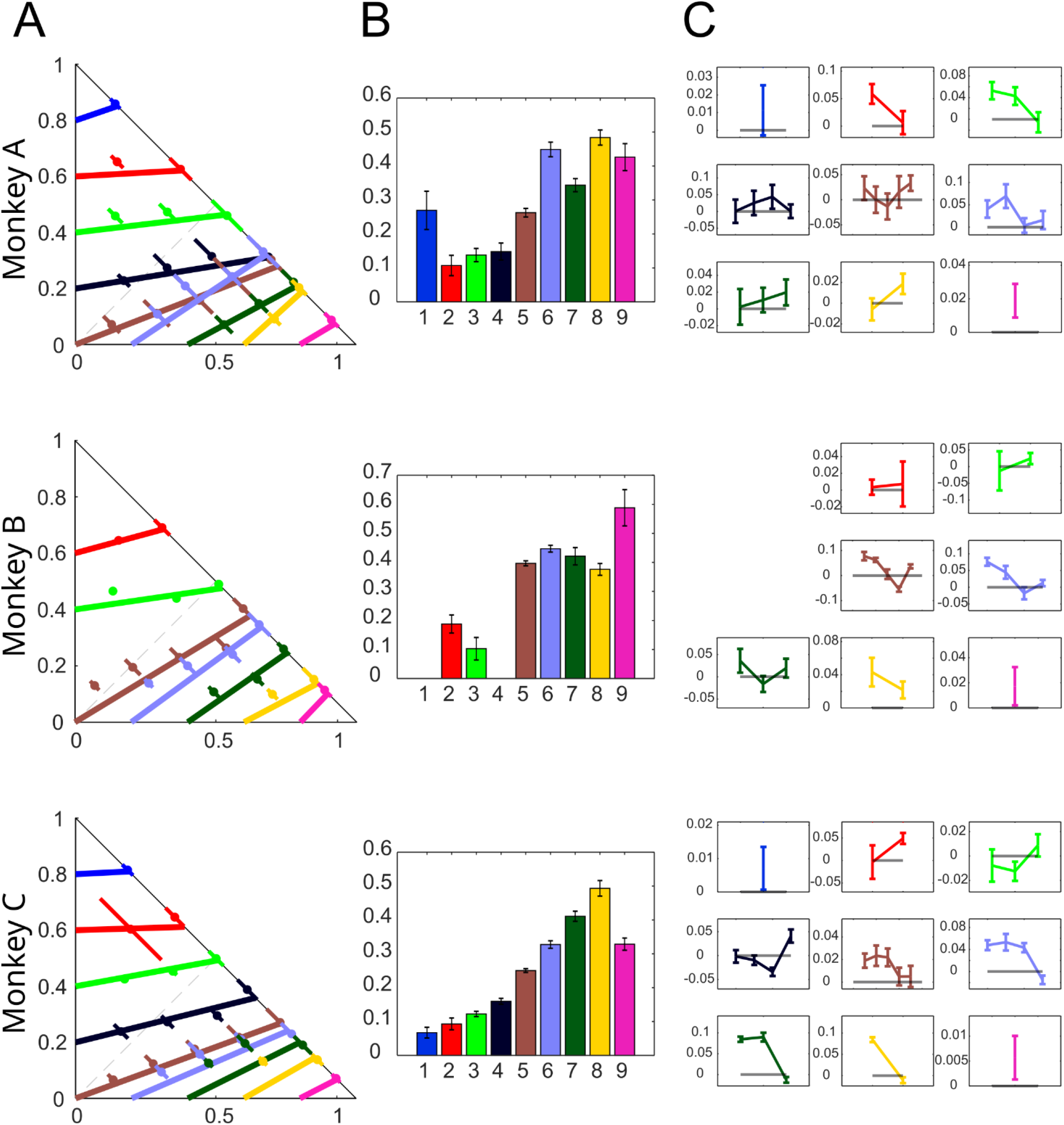
Out-of-sample test for linearity and parallelism of indifference curves in the Marschak-Machina triangle. (A) Indifference curves estimated with linear least-squares and out-of-sample indifference points. Dots represent the mean of indifference points across all sessions (that fall within the same indifference curve with the same color); lines show Standard Deviation (SD) of IPs across all sessions. (B) Bar graphs showing significant differences in slope of the indifference curves estimated with linear least-squares (mean ± SEM; one-way ANOVA p < 0.001 for all three animals). The nine colors correspond to the nine indifference curves being tested. (C) Line plots showing significant residuals between indifference points and estimated indifference curves (mean ± SEM; p<0.05; one-sample t-test against indifference curves, represented by grey lines). The y-axis represents the residuals (0 = same as estimated indifference curves), and the x-axis represents the position of the indifference points in the Marschak-Machina triangle.

